# Single cell spatial transcriptomics identifies coordinated cellular programs associated with good prognosis in microsatellite stable colorectal cancer

**DOI:** 10.1101/2025.03.31.646286

**Authors:** Victoria T. Karlsen, Umair Majid, Håvard T. Lindholm, Sinan U. Umu, Karoline Rapp Vander-Elst, Ann-Christin Røberg Beitnes, Marianne A. Merok, Kjersti Thorvaldsen Hagen, Hogne Røed Nilsen, Sheraz Yaqub, Espen S. Bækkevold, Diana Domanska, Frode L. Jahnsen

## Abstract

It has been reported that microsatellite instable (MSI) tumors in colorectal cancer (CRC) exhibit stronger anti-tumor responses than microsatellite stable (MSS) tumors. Analyzing scRNA-seq data from 185 CRC patients we found that immune, structural, and cancer cells in MSS tumors with high numbers of tumor-infiltrating CD8 T cells and macrophages (TAMs; CD8^hi^TAM^hi^) and CD8^low^TAM^hi^ tumors were enriched for pathways associated with anti- and pro-tumor responses, respectively. In CD8^hi^TAM^hi^ tumors, TAMs expressed an IFN-induced phenotype (e.g. *GBP1, CXCL9, IFITM3*) and high infiltration of *GBP1*+ TAMs was associated with better overall survival (n=941). High-resolution spatial transcriptomics (Visium HD) revealed that *GBP1*+ TAMs clustered with *CXCL13*+*IFNG+PDCD1+* CD8 T cells in tumoral niches, suggesting that *GBP1*+ TAMs were involved in recruitment and activation of tumor-reactive CD8 T cells. Together, we uncovered coordinated cellular programs across cell types in the microenvironment of MSS tumors, based on simple classification variables, that were strongly associated with prognosis.

## INTRODUCTION

Colorectal cancer (CRC) is one of the most common cancer types world-wide with a 5-year overall survival rate of 60-70% ^1–3^. CRC consists of two genetically distinct subtypes. The majority is microsatellite stable/mismatch repair proficient (MSS/MMRp) tumors constituting approximately 85% of cases, whereas up to 15% are microsatellite instable/mismatch repair deficient (MSI/MMRd) tumors.

The latter has high mutational burden, whereas MSS tumors have low mutational burden^4,5^. Furthermore, MSI tumors display more anti-tumor immunity than MSS tumors^6–8^, and MSI tumors respond strongly to immune checkpoint blockade (ICB)^9–11^. ICB has therefore recently been introduced as neoadjuvant treatment to CRC patients with locally advanced MSI tumors^9^. MSS tumors have been believed to be unresponsive to ICB, but recently it has been shown that a fraction of patients with MSS tumors respond well to this treatment^10,11^. Moreover, single-cell transcriptomic analysis of CRC patients showed that a subgroup of MSS tumors were transcriptionally more similar to MSI tumors than other MSS tumors^12^. In line with this, genomic analysis CRC cells has shown that MSS tumors consist of distinct clusters that are related to prognosis^13^. Together, these findings indicate that MSS tumors are heterogeneous both related to their genomic landscape, transcriptional phenotype, and response to ICB.

It is widely recognized that high infiltration of T cells is linked to a more favorable prognosis in various cancer types including CRC^14^. In contrast, infiltration of tumor-associated macrophages (TAMs), on the other hand, has been associated with worse prognosis in many cancer types^14–16^, suggesting that TAMs may play a role in promoting tumor progression. However, reports on the prognostic effect of TAMs in CRC have shown contradictory results^17–19^. We recently showed that the prognostic effect of TAM infiltration in MSS CRC highly depended on the concurrent number of infiltrating T cells^20^. The best clinical outcome was found when patients had MSS tumors with coexisting high numbers of TAMs and T cells, whereas MSS tumors with high numbers of TAMs and low T cell counts had the worst prognosis^20^. As we know that resident colonic macrophages consist of many functionally distinct subtypes^21^ our findings suggest that the composition of TAM subsets/cell states in TAM-rich tumors differ depending on the number of tumor-infiltrating T cells. To study TAMs in CRC in more detail and explore their functional roles within the complex tumor microenvironment (TME) we created a high-resolution single cell RNA-sequencing (scRNA-seq) atlas of CRC encompassing 185 patients and a total of 890.889 cells, representing 68 annotated cell subsets. The scRNA-seq data was integrated with high resolution spatial transcriptomics to map the spatial organization of all cell types within the TME, enabling the identification of cellular neighborhoods and their receptor-ligand interactions. Leveraging this fine-grained spatial atlas, we explored the cellular and molecular mechanisms underlying the prognostic significance of TAM and CD8 T cell densities.

## RESULTS

We integrated and harmonized cells from tumor and adjacent normal colon tissue profiled by scRNA-seq data from seven published^8,12,22–26^ and one unpublished data set with a total of 185 patients. Clinical characteristics are given in Supplementary Figure 1A. All samples were concatenated into a single object, prior to performing quality control where low-quality cells were removed. Integration of the concatenated object was performed using single-cell variational inference (scVI). A total of 890.889 high quality cells were included.

The final integrated data was annotated into 12 major cell lineages (Supplementary Figure 1B). The relative abundance of the cell types varied between the patients (Supplementary Figure 1C). The cell lineages were further sub-clustered into 68 cell types (Supplementary Figure 1B) that were manually annotated using published data based on top differentially expressed genes (DEGs, Supplementary Table 1). Automatic annotation such as CellTypist models were tested, but since most such models are based on cells from healthy tissue they did not fit well with tumor-associated cell types. Moreover, to get a deeper understanding of the functional roles of cells in the complex TME, we combined scRNA-seq data and high-resolution spatial transcriptomics using Visium HD (10X Genomics). This is a probe-based platform that identifies approximately 19.000 genes combined with hematoxylin and eosin (H&E) staining on the same tissue section.

### High resolution spatial transcriptomic analysis of cancer cells in CRC

Integrated scRNA-seq analysis of epithelial cells from tumor and normal colonic tissue divided them into 11 clusters (Figure 1A). For differential abundance testing we used Milo^27^ which showed that four clusters were more abundant in the tumor tissue (Figure 1B). These cell clusters expressed an intestinal stem cell (Stem)/transit-amplifying (TA)-like phenotype (*LGR5, ASCL2*), whereas clusters that were more abundant in normal colon included differentiated epithelial cells such as goblet cells (*MUC2, MUC4*), colonocytes (*CA2*), and tuft cells (*LRMP*) (Figure 1C).

**Figure 1:**
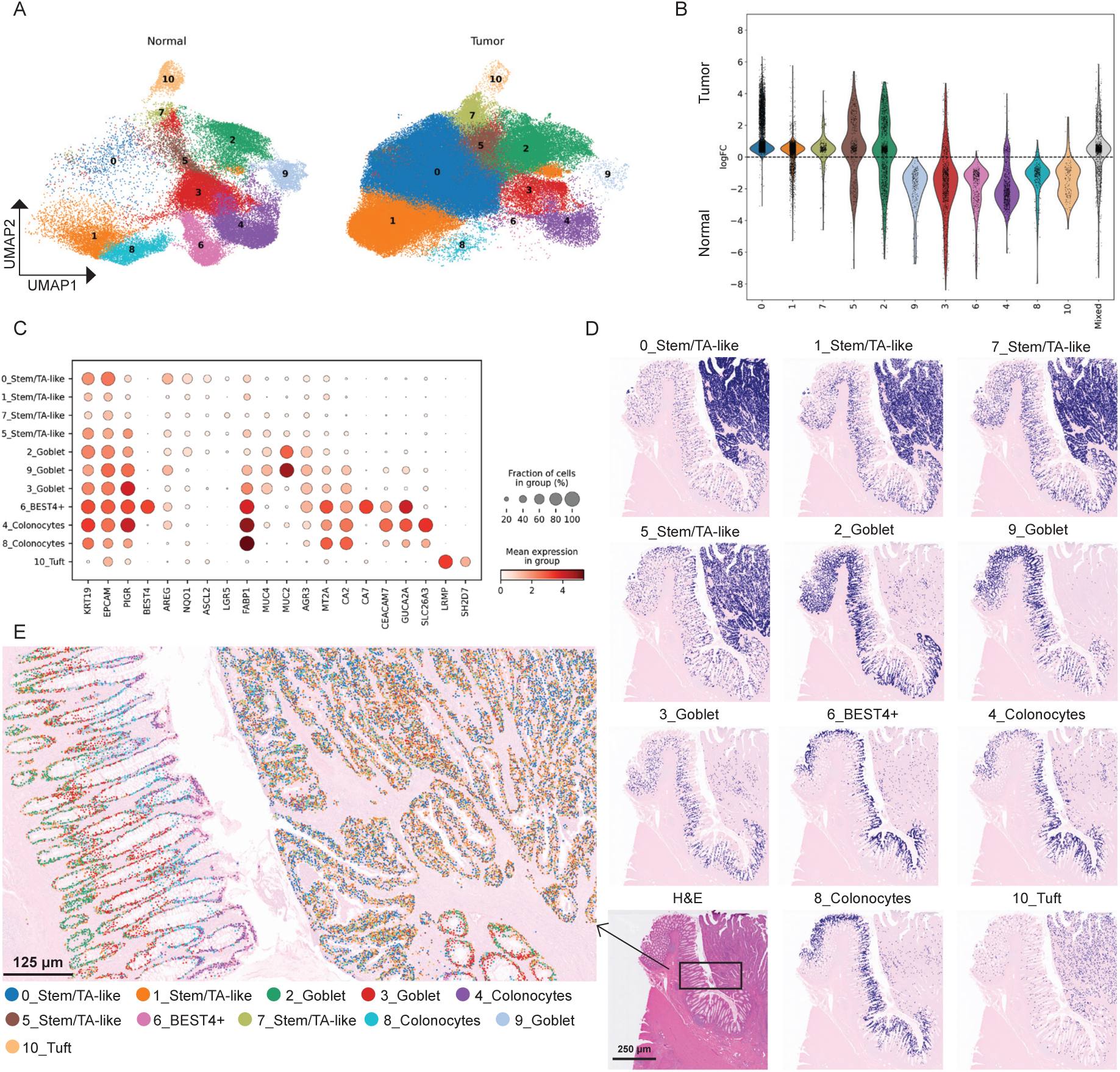
Analysis of epithelial cells with scRNA-seq and Visium HD data. **A)** UMAPs with fine-grained clustering of epithelial/tumor cells from all normal colon (left) and tumor tissue samples (right). **B)** Differential abundance analysis (Milo) comparing the abundance of epithelial cell types from normal colon and tumor tissue. **C)** Marker gene dot plot for epithelial cell types from normal colon and tumor tissue with predicted annotations indicated (left). **D)** Spatial visualization of the annotated epithelial/tumor clusters (A-C) combined with Visium HD (Easydecon pipeline) on FFPE tissue section from CRC (Sample P5^83^; each cell type is shown in separate images for simplicity). H&E staining (lower left). **E)** High magnification image showing the spatial distribution of all cell clusters separated by different colors (Bin2Cell, Easydecon). Note the well-defined organization of differentiating epithelial cells from crypt bottom to crypt top (left) in contrast to random distribution of mainly undifferentiated epithelial cells in the tumor tissue (right). Compiled data for all tumor samples are shown in Supplementary Figure 2.

To validate these findings in situ, we combined the annotated scRNA-seq data and Visium HD spatial gene expression on sections of formalin-fixed paraffin-embedded (FFPE) tumor tissue from CRC patients (n=7; clinical characteristics in Supplementary Table 2). Easydecon, our recently developed pipeline^28^, was used to transpose cell types identified with scRNA-seq in Visium HD data. By combining H&E staining and visualization of annotated cell types on the same tissue sections we found that clusters dominated by Stem/TA-like phenotypes were mainly expressed in tumor cells, whereas genes that marked differentiated epithelial cells were mainly expressed in non-tumor epithelium (Figure 1D, Supplementary Figure 2).

Our analysis using Easydecon is based on the default resolution of 8×8 µm bin size. To examine the spatial transcriptome at the single cell level we took advantage of the recently published pipeline Bin2cell^29^ that reconstructs cells by leveraging morphology image segmentation and gene expression information at the highest resolution (2μm bins). Combining Bin2cell and Easydecon the normal epithelial differentiation from crypt bottom to crypt top was readily appreciated at the single cell level in the normal colonic mucosa, whereas cancer cells co-expressed absorptive and secretory gene programs together with Stem/TA-like gene programs, reflecting substantial dysregulated gene expression in tumor cells (Figure 1E).

To determine how Visium HD performed as a stand-alone platform we manually annotated the tissue regions that contained cancer cells and normal epithelial cells on H&E-stained images of three samples that contained both tissues (Supplementary Figure 3A). Clustering of all bins within the annotated areas resulted in 30 clusters (Supplementary Figure 3B), in which nine expressed high levels of the pan-epithelial marker *EPCAM* (Supplementary Figure 3C). The six major clusters derived from cancer-annotated areas all expressed a stem cell (Stem)/transit-amplifying (TA)-like phenotype (*LGR5, ASCL2*), whereas the clusters derived from normal epithelium expressed high levels of genes typical for absorptive and secretory cells (*PIGR, MUC2, GUCA2A*) (Supplementary Figure 3D). Based on this finding, the gene profiles expressed in the six cancer-cell specific clusters were used to identify tumor cells to determine their spatial relationship to other cell types within the TME in the following sections.

### Tumor-associated macrophage (TAM) subsets are organized in specific tumor niches

Integrated analysis of scRNA-seq data from normal colon and tumor tissue divided macrophages into 11 clusters (Figure 2A). Most clusters were more abundant in tumor tissue (Figure 2B). *SPP1+* TAMs (*SPP1, APOE, ACP5, APOC1, TREM2*) associated with tumor-promoting properties^24,30,31^, *GBP1+* TAMs (*GBP1, CXCL9, CXCL10, IDO1*) associated with anti-tumor properties^32,33^, and *HILPDA+* TAMs (expressing *HILPDA, CXCL8, CCL20, INHBA* and low levels of *CD163*) were mainly found in tumor tissue (Figure 2A-C). Recently elicited monocytes, designated *S100A8+* TAMs (*S100A8, S100A9*), proinflammatory *IL1B+* TAMs, and *FOLR2+* TAMs expressing a tissue resident phenotype (*FOLR2, LYVE1, SELENOP, MRC1*/CD206*, F13A1*) were also more abundant in tumor tissue, but these clusters were also present in normal tissue (Figure 2A-C). Proliferating (*MKI67+*) TAMs were only found in tumor tissue, expressing a phenotype very similar to *SPP1+* TAMs (Figure 2A-C). Two subsets expressed high levels of HLA class II genes. The largest cluster was more abundant in tumor tissue (HLA class II^hi^ TAMs), whereas the other cluster was virtually only found in normal tissue. Interestingly, the latter subset expressed *DNASE1L3* and *ADAMDEC1* suggesting an embryonic origin^21^. The final cluster expressed both macrophages and dendritic cell genes compatible with cDC3^21^ (Figure 2C).

**Figure 2:**
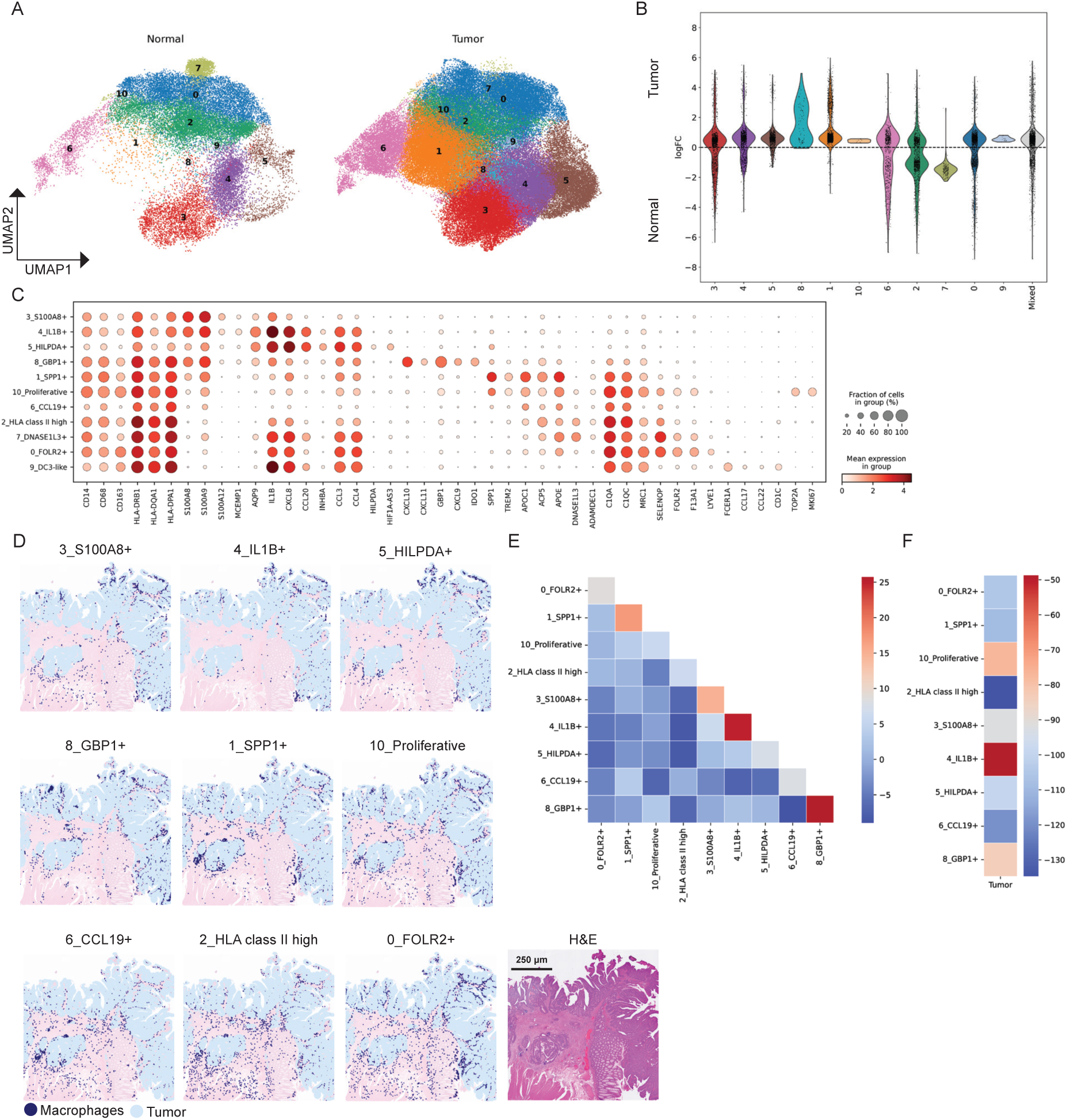
Analysis of macrophages with scRNA-seq and Visium HD data. **A)** UMAPs with fine-grained clustering of macrophages from all normal colon (left) and tumor tissue samples (right). **B)** Differential abundance analysis (Milo) comparing the abundance of macrophage cell types from normal colon and tumor tissue. **C)** Marker gene dot plot for macrophage cell types from normal colon and tumor tissue with predicted annotations indicated. **D)** Spatial visualization of annotated macrophage cell types (A-C) combined with Visium HD (Easydecon) on FFPE tissue section from CRC (Sample P2^83^; each cell type is shown in separate images for simplicity). H&E staining (lower right). **E)** Neighborhood enrichment analysis (Squidpy) of TAM cell types on CRC tissue sections (n=7). **F)** Neighborhood enrichment analysis (Squidpy) examining the average distance between TAM cell types and tumor cells on CRC tissue sections (n=7).

Visium HD showed that TAM subsets were densely located at the tumor front and in fibrous strands between the tumor cells (Figure 2D, Supplementary Figure 2). To perform a more detailed unbiased spatial analysis, we utilized Squidpy^34^, a pipeline for identifying spatial patterns in spatial transcriptomics data. It showed that some TAM subtypes were highly co-enriched with themselves, in particular *SPP1*+TAMs, *S100A8*+ TAMs, *IL1B*+ TAMs and *GBP1*+ TAMs (Figure 2E). *IL1B*+TAMs, proliferative TAMs, and GBP1+ TAMs were located closest to tumor cells (Figure 2F).

Together, this showed that TAMs consisted of many functionally distinct subsets, of which some were spatially organized in distinct tumor niches.

### Extensive reprogramming of T cells in tumor tissue

T cells, natural killer (NK) cells, and innate lymphoid cells (ILCs) were divided into 11 clusters (Figure 3A). Using the Milo pipeline, we found that most subsets were more abundant in tumor tissue compared to their normal counterpart (Figure 3B). The CD4 T cell compartment contained five subsets. *CXCL13*-expressing CD4 T cells were virtually only present in tumor tissue and expressed low levels of both immune activation genes (*IFNG, CD40LG*) and inhibitory genes (*PDCD1, CTLA4, HAVCR2*, *LAG3*) (Figure 3C), resembling a tumor-reactive phenotype identified in many cancer settings^35–38^. Enrichment of this phenotype has also been associated with response to ICB^35^.

**Figure 3:**
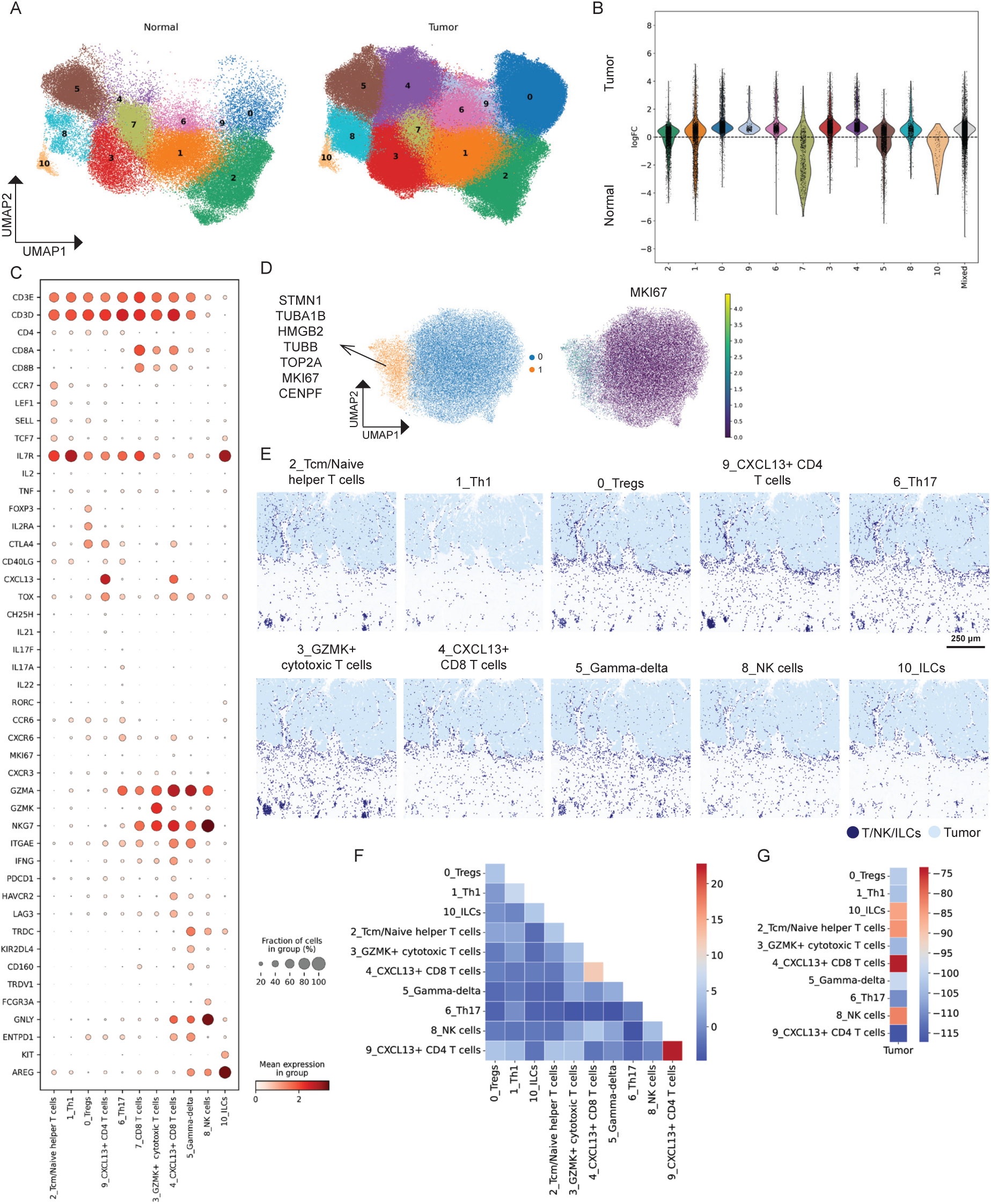
Analysis of T cells, NK cells, and ILCs with scRNA-seq and Visium HD data. **A)** UMAPs with fine-grained clustering of T cells, NK cells, and ILCs from all normal colon (left) and tumor tissue (right). **B)** Differential abundance analysis (Milo) comparing the abundance of T cell, NK cell, and ILC types from normal colon and tumor tissue. **C)** Marker gene dot plot for T cell, NK cell, and ILC types from normal colon and tumor tissue with predicted annotations indicated. **D)** UMAP of *CXCL13+* CD8 T cells (from tumor only) showing that a subset (9%) expressed several proliferation markers (left). Feature plot showing the expression of *MKI67* (right). **E)** Spatial visualization of annotated T cell, NK cell, and ILC types (A-C) combined with Visium HD (Easydecon) on FFPE tissue section from CRC (each cell type is shown in separate images for simplicity). **F)** Neighborhood enrichment analysis (Squidpy) examining the average distance of T cells, NK cells, and ILC types on CRC tissue sections (n=7). **G)** Neighborhood enrichment analysis (Squidpy) examining the average distance between T cell, NK cell, and ILC types with tumor cells on CRC tissue sections (n=7).

The CD4 T cell compartment also included Tregs (*Foxp3, CTLA4, IL2RA*), Th1-like cells (*TNF, IL2, CD40LG, GZMA, NKG7*), Th17-like cells (*IL17, RORC, IL22, GZMA*), and T cells with a naïve/central memory phenotype (*CCR7, SELL, LEF1, TCF7*) (Figure 3C).

The CD8 T cell compartment contained a *CXCL13*-expressing subset that was virtually only identified in tumor tissue (Figure 3A, B). *CXCL13*+ CD8 T cells expressed activation genes (*IFNG, GZMA, NKG7, GNLY*) as well as inhibitory genes (*PDCD1, HAVCR2, LAG3, ENTPD1*) (Figure 3C); a phenotype designated exhausted tumor-reactive CD8 T cells in several cancer settings^39,40^. However, as previously reported in CRC, a significant fraction (9%) of tumor-infiltrating *CXCL13*+ CD8 T cells expressed proliferative markers including *MKI67* (Figure 3D) as well as the highest levels *IFNG* and other immune activation genes (Figure 3C), indicating that they had not lost their anti-tumor effector potential^41^.

The other CD8 T cell subsets were *CXCL13*-negative. One subset was more frequent in the tumor tissue and expressed an effector phenotype with significant levels of *GZMK, NGK7, GZMA,* and *IFNG,* and low levels of inhibitory genes (*PDCD1, HAVCR2, LAG3*) (Figure 3A-C). *GZMK-*expressing tumor-specific CD8 effector T cells have been found in several cancer types^35,42^ and have been associated with response to ICB^35^. The other subset of *CXCL13*-CD8 T cells was more abundant in normal tissue and expressed a resident memory phenotype (*ITGAE, CD160, IFNG, GZMA, NKG7*) (Figure 3A-C).

𝛾/𝛿 T cells (I*TGAE, TRDC, KIR2DL4, CD160*) and ILCs (*KIT, AREG*) were found equally abundant in tumor and normal tissue, whereas NK cells (*FCGR3A, GNLY*) were more abundant in tumor tissue (Figure 3A-C).

Visium HD showed that the clusters were located both at the tumor border and to a variable extent infiltrating tumor tissue (Figure 3D, Supplementary Figure 2). *CXCL13*+ CD4 T cell subset was strongly co-enriched with itself, followed by *CXCL13*+ CD8 T cells (Figure 3F). The other subsets showed little tendency to cluster (Figure 3F). *CXCL13*+ CD8 T cells were spatially closest to tumor cells (Figure 3G).

Together, we found that the phenotype of tumor-associated T cells was very different from their normal counterparts, and some of these subsets were spatially organized in specific niches within the tumor tissue.

### T cells show spatial proximity to myeloid cells in tumor tissue

Studies examining the effect of ICB have shown that T cell clones that expand on therapy are found within the tumor pre-treatment, suggesting that tumor-reactive T cells are activated locally by antigen presenting cells (APCs)^35^. To explore this, we first analyzed the dendritic cell compartment. Dendritic cells included cDC1, cDC2, cDC3 and mregDCs (mature DCs enriched in immunoregulatory molecules)^41,43^. mregDCs were virtually only found in tumor tissue, whereas the other DC subsets were situated in both tumor and normal colon (Figure 4A, B). Spatial visualization showed that mregDCs clustered together whereas the other DCs subsets were distributed both in the tumor stroma and intratumorally (Figure 4D, Supplementary Figure 2). We also detected low numbers of plasmacytoid DCs in tumor tissue (not shown).

**Figure 4:**
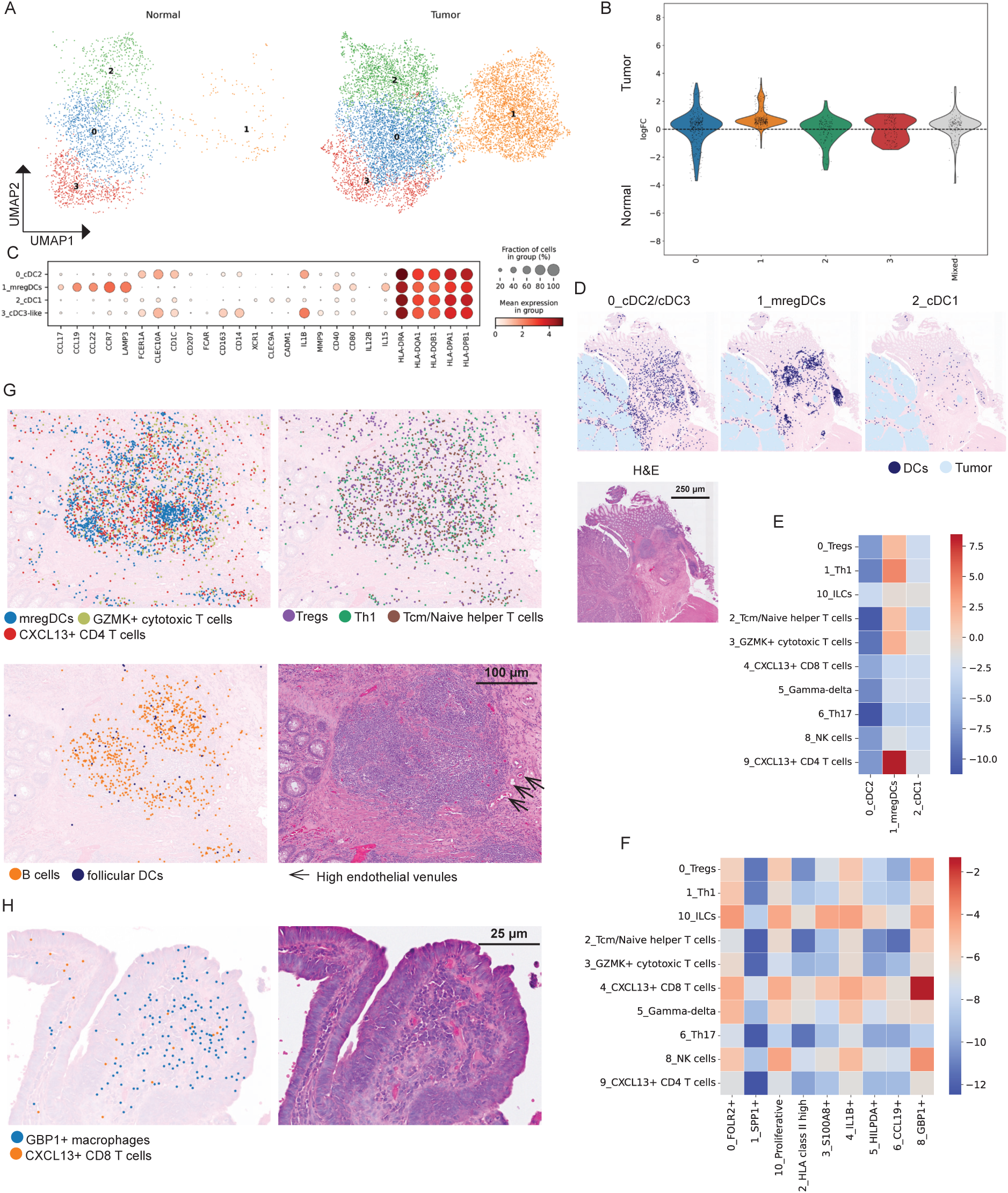
Spatial analysis of tumor-infiltrating T cells and myeloid cells. **A)** UMAPs of DC subsets from all normal colon (left) and tumor tissue (right). **B)** Differential abundance analysis (Milo) comparing the abundance of DC subsets from normal colon and tumor tissue. **C)** Marker gene dot plot for DC subsets from normal colon and tumor tissue with predicted annotations indicated. **D)** Spatial visualization (Easydecon) of annotated DC subsets (A-C) combined with Visium HD on FFPE tissue section from CRC (Sample P1^83^; each cell type is shown in separate images for simplicity). H&E staining (bottom). **E)** Neighborhood enrichment analysis (Squidpy) examining the average distance between T cell/NK cell/ILC types and DC subsets on CRC tissue sections (n=7). **F)** Neighborhood enrichment analysis (Squidpy) examining the average distance between T cells/NK cells/ILC types and TAM subsets on CRC tissue sections (n=7). **G)** Spatial visualization (Sample P1^83^; Bin2Cell, Easydecon) of mregDCs, *GZMK+* CD8 T cells, and *CXCL13*+ CD4 T cells (upper left); Tregs, naïve/central memory Th cells, and Th1 cells (upper right); follicular DCs and B cells (lower left); and H&E staining of TLS from the same section of tumor tissue (Sample P1^83^). Arrows indicate high endothelial venules (HEVs). Scare bar 100 µm. **H)** High magnification image showing colocalization of *GBP1*+ TAMs and *CXCL13*+ CD8 T cells intratumorally (Sample P2^83^; Bin2Cell, Easydecon). Scare bar 25µm.

Neighborhood enrichment analysis showed that, among the DC subsets, mregDCs were close to CXCL13+ CD4 T cells, Th1 cells, Tregs, naïve/cm T cells, and *GZMK*+ CD8 T cells (Figure 4E) and spatial visualization revealed that mregDCs and the T cell subsets were organized in lymphoid aggregates (Figure 4G). Interestingly, the proximity of mregDCs, *CXCL13*+ CD4 T cells, and *GZMK*+ CD8 T effector cells has been shown to correlate with response to ICB^35^. mregDCs expressed high levels of activation genes such as HLA class II genes, *CD40, CD80, CD86* and *PDCD1LG2* (PDL2), *IL12B* important for Th1 differentiation, and IL15 involved in CD8 T cell activation^44^ (Figure 4C). They also expressed the chemokines *CCL19*, *CCL17* and *CCL22* that attract both naive and memory T cells^35,43^. B cells were also localized in these lymphoid aggregates together with follicular dendritic cells (*CR1*, *CR2*; Figure 4G), and high endothelial venules (Figure 4G), demonstrating that many of these aggregates were mature tertiary lymphoid structures (TLS).

TAMs have also been shown to be involved in the activation of tumor-reactive T cells^45,46^. Examining the spatial proximity between TAMs and T cells, we found that the highest enrichment score was between *CXCL13*+ CD8 T cells and *GBP1*+ TAMs (Figure 4F). Spatial visualization showed that these cell types were organized within distinct niches intratumorally (Figure 4H). *GBP1*+ TAMs expressed high levels of *CXCL9*, *CXCL10* and *CXCL11 (*Figure 2C); all ligands for the receptor CXCR3 expressed on *CXCL13*+ CD8+ T cells (Figure 3C). Together, this suggested that *CXCL13*+ CD8 T cells were attracted and activated by *GBP1*+ TAMs.

Interestingly, *SPP1+* TAMs showed the lowest enrichment score with all T cell subsets (Figure 4F). This is in line with previous results showing that *SPP1+* TAMs limit T cell infiltration and thus promote tumor progression^47^.

### B cells accumulate in TLS whereas IgG+ plasma cells are most prominent in the tumor stroma

The B cell compartment contained naïve (*IGHD*), germinal center (*MKI67, AICDA, BCL6*) and memory B cells (Supplementary Figure 4A, C). Differential abundance analysis showed that they were similarly present in tumor and normal tissue and spatial visualization showed that B cells were virtually confined to TLS in the tumor stroma and in isolated lymphoid follicles (ILFs) in normal colon (Supplementary Figures 2, 4B, 4D, Figure 4G). Plasma cells were divided into three clusters. One cluster expressed high levels of *IGHA1* and *IGHA2*; the second cluster expressed *IGHG1*-*IGHG4* subclasses, and the third cluster expressed high levels of IGHM (Supplementary Figure 4E, G). Visium HD showed variable levels of plasma cells in the tumor stroma; mainly expressing *IgHG*, whereas normal colonic tissue was dominated by *IGHA* and a small number of *IGHM* (Supplementary Figure 2 and 4F, H).

### Stromal cells are a prominent component of the tumor microenvironment

Fibroblasts were divided into six clusters; two were virtually only present in the tumor tissue (Figure 5A, B). These two cancer-associated fibroblasts (CAFs) expressed genes such as *MMP1*, *MMP3*, *CXCL8*, *CTHRC1*, and *LRRC15* (Figure 5C). Genes that have been associated with both inflammatory and immunosuppressive functions in different cancer and inflammatory settings^48–51^. Both CAF subsets expressed a contractile program including *ACTA2* and *TAGLN*, suggesting myofibroblast differentiation.

**Figure 5:**
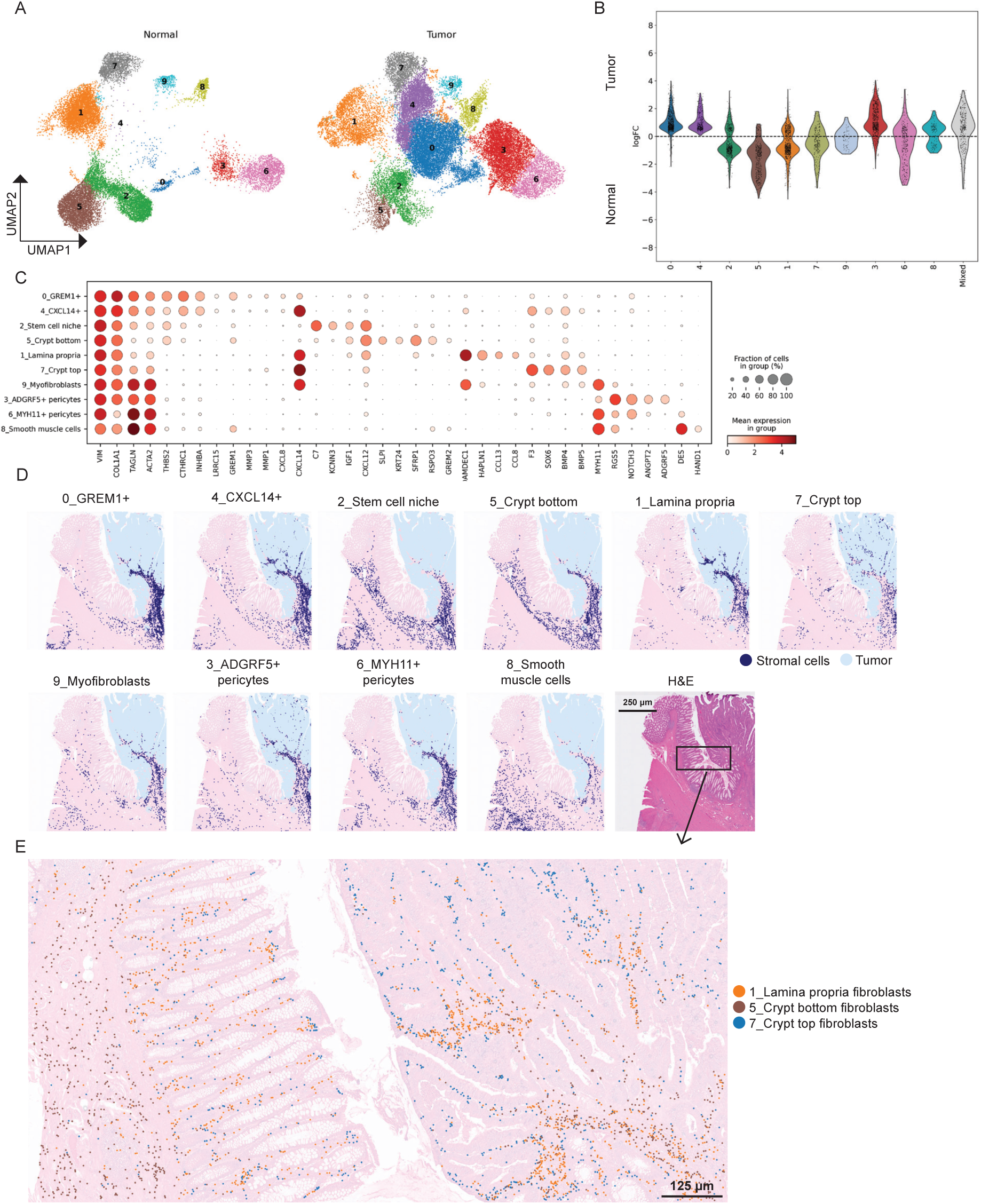
Analysis of stromal cells with scRNA-seq and Visium HD data. **A)** UMAPs with fine-grained clustering of stromal cell types from normal colon (left) and tumor tissue (right). **B)** Differential abundance analysis (Milo) comparing the abundance of stromal cell types from normal colon and tumor tissue. **C)** Marker gene dot plot for stromal cell types from normal colon and tumor tissue with predicted annotations indicated. **D)** Spatial visualization of the annotated stromal cell clusters (A-C) combined with Visium HD (Easydecon) on FFPE tissue section from CRC (Sample P5^83^; each cell type is shown in separate images for simplicity). H&E staining (lower right). **E)** High magnification image showing the spatial distribution of selected stromal cell clusters separated by different colors (Sample P5^83^; Bin2Cell, Easydecon). Note the well-defined positioning of CT, LP, CB fibroblasts in normal colonic mucosa (left) and similar organization of their tumor counterparts (right).

Spatial visualization showed that these subsets were located in the tumor stroma close to the tumor front (Figure 5D, Supplementary Figure 2). The other four fibroblast subsets were more abundant in normal colonic tissue (Figure 5B) and their transcriptomic profiles were in agreement with previous reports studying the function of fibroblasts related to epithelial differentiation^52^. This included crypt top (CT) fibroblasts (*F3, SOX6, BMP4, BMP5, CXCL14*), lamina propria (LP) fibroblasts (*ADAMDEC1, HAPLN1, CCL13, CXCL14*), crypt bottom (CB) fibroblasts (*IGF1, SFRP1, GREM1, RSPO3*), and stem cell niche (SCN) fibroblasts (*C7*)^52^ (Figure 5C). Spatial visualization showed that fibroblasts in the normal colonic mucosa were positioned along the crypt bottom-top axis according to their phenotype (Figure 5E). Interestingly, their phenotypic counterparts residing in the tumor tissue showed very similar spatial distribution (Figure 5E), suggesting that these cells contributed to tumor growth as suggested by others^8^.

Pericytes were divided into two subsets (Figure 5A, C). *ADGRF5*+ pericytes were more abundant in tumor tissue (Figure 5B). These pericytes expressed *ANGPT2* that promotes angiogenesis^53^. *MYH11*+ pericytes, on the other hand, were mostly found in the tumor stroma and the normal colonic tissue (Figure 5D and Supplementary Figure 2).

### Endothelial tip cells and activated post-capillary venules are increased in tumor tissue

The endothelial cell compartment contained 10 clusters including capillary, venous, arterial, and lymphatic endothelial cells (Figure 6A, C). Three clusters of capillary endothelial cells were highly abundant in tumor tissue, which included Tip cells (*ESM1, CXCR4, PGF, FOLH1, ANGPT2*) (Figure 6B, C). ESM-1 has been identified as a specific biomarker of tip cells during neoangiogenesis^54^. Two clusters of capillary endothelial cells, expressing *CD36*, were more abundant in normal tissue (Figure 6B). Venous endothelial cells (*ACKR1*) consisted of three subsets; two were mainly tumor-derived (Figure 6B). The venous vessels were located mainly in the tumor stroma (Figure 6D and Supplementary Figure 2) and expressed several genes encoding adhesion molecules involved in leukocyte trafficking (e.g. *SELP*, *SELE*, *VCAM*-1 and *MADCAM-1*) (Figure 6C), suggesting that they are important for recruitment of leukocytes to the tumor site. Lymphatic endothelial cells (C*CL21, LYVE1, TFF3*) and arterial endothelial cells (*FBLN5, GJA5*) were mainly located in the tumor stroma as well as in the normal the colonic mucosa^55^ (Figure 6D).

**Figure 6:**
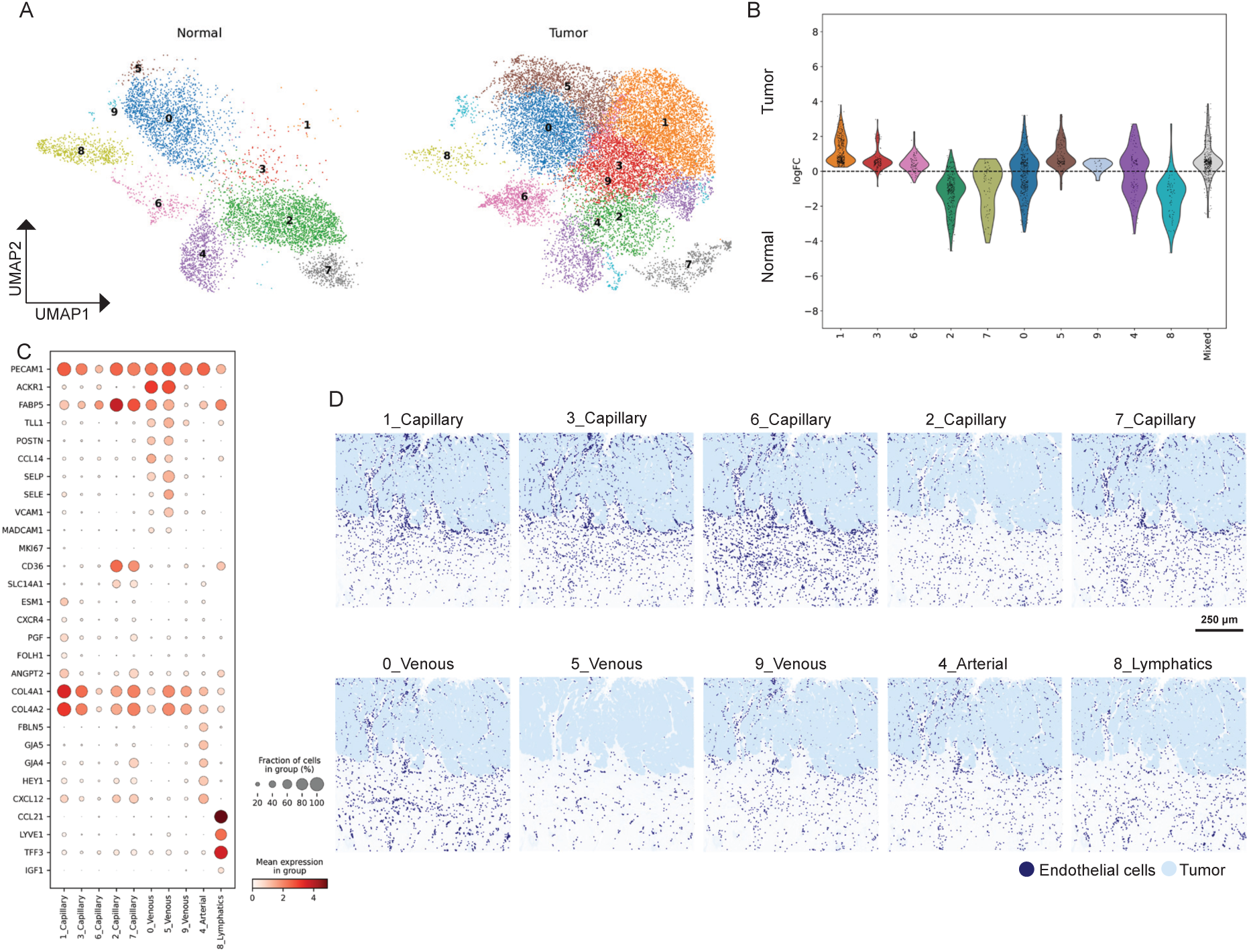
Analysis of endothelial cells with scRNA-seq and Visium HD data. **A)** UMAPs of fine-grained clustering of endothelial cell types from normal colon (left) and tumor tissue (right). **B)** Differential abundance analysis (Milo) comparing the abundance of endothelial cell types from normal colon and tumor tissue. **C)** Marker gene dot plot for endothelial cell types from normal colon and tumor tissue with predicted annotations indicated. **D)** Spatial visualization of the annotated endothelial cell clusters (A-C) combined with Visium HD (Easycon) on FFPE tissue section from CRC (each cell type is shown in separate images for simplicity).

### Reprogramming of TAMs is associated with better prognosis in MSS tumors

By examining the prognostic effect of infiltrating T cells and TAMs in CRC, we recently found that concomitant high numbers of T cells and TAMs were associated with good prognosis in patients with MSS tumors, whereas MSS tumors with low numbers of T cells and high numbers of TAMs had by far the worst prognosis^20^. Based on this finding we hypothesized that TAMs in the two situations were functionally different. To explore the underlying mechanisms for this difference we stratified the CRC patients into four equal groups depending on the density of CD8 T cells and TAMs (Figure 7A) and performed a comparative transcriptome analysis of MSS patients with CD8^hi^TAM^hi^ tumors (n=14; 14% of MSS tumors) and CD8^low^TAM^hi^ tumors (n=29; 29% of MSS tumors). Using Milo, we found that three TAM clusters, *SPP1*+ TAMs, *FOLR2*+ TAMs, and class II^hi^ TAMs, were more abundant in the CD8^low^TAM^hi^ group (Figure 7B). To investigate whether the prognostic difference between the groups was caused by a change in the relative representation of TAM subsets or whether the TAMs were transcriptionally reprogrammed across the subtypes, we determined the biological pathways in all TAMs based on DEGs between the two groups. We found that TAMs in CD8^hi^TAM^hi^ tumors were enriched for pathways that are associated with anti-tumor responses, including antigen processing and presentation/cross presentation, as well as interferon (IFN) type I and II signaling (Figure 7C)^56^, whereas TAMs in CD8^low^TAM^hi^ tumors were enriched for pathways which have been associated with tumor-promoting activities, such as TGF-b signaling, VEGFA-VEGFR2 pathway, and PDGF signaling (Figure 7C)^56–58^.

**Figure 7:**
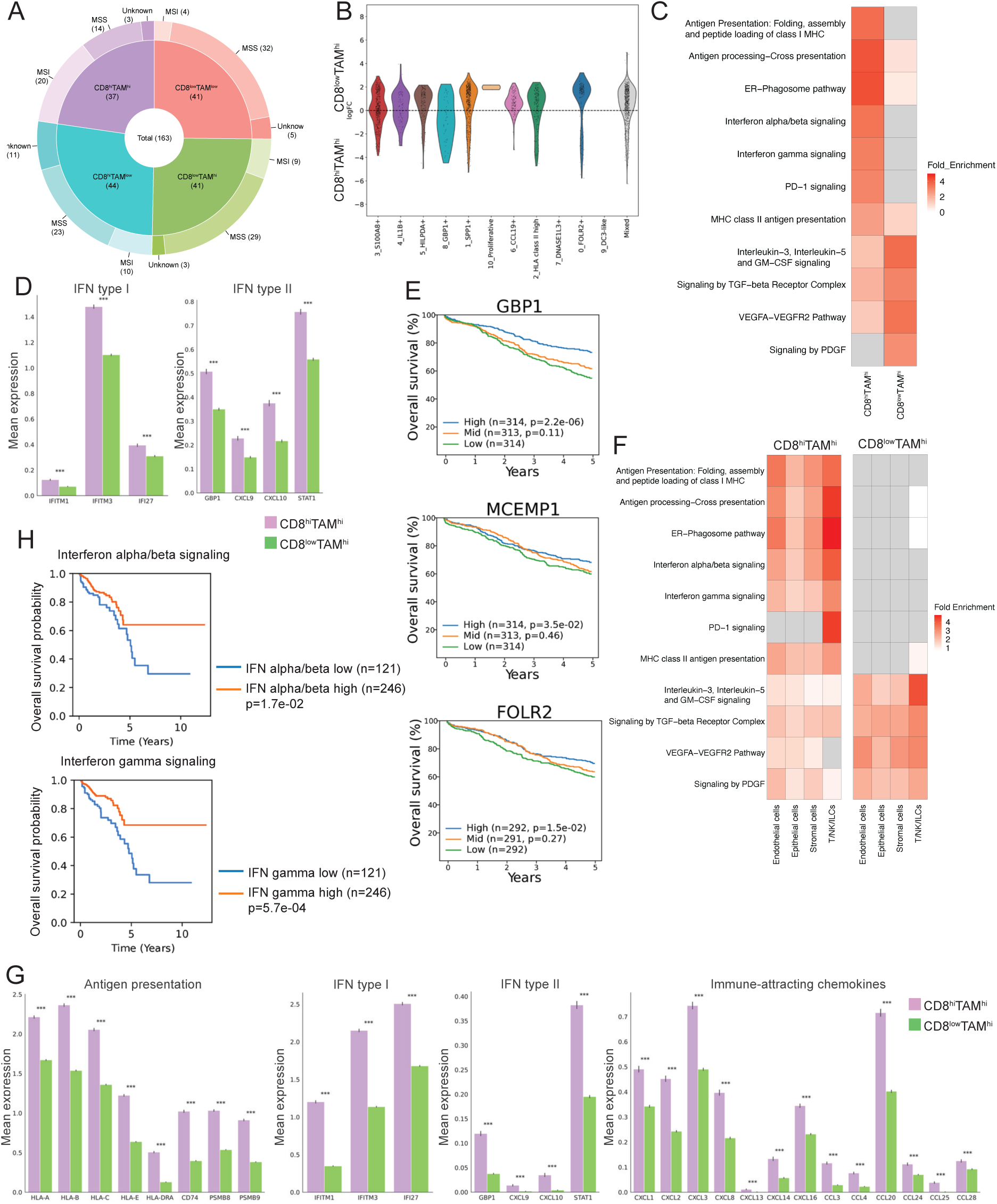
Transcriptomic analysis comparing TAMs, structural cells and immune cells in CD8^low^TAM^hi^ and CD8^hi^TAM^hi^ MSS tumors. **A)** Pie-donut plot dividing CRC patients in four equally-sized groups depending on whether the tumor has high or low numbers of infiltrating CD8 T cells and TAMs. The numbers of patients with MSS and MSI tumors in each group are indicated in brackets. **B)** Differential abundance analysis (Milo) comparing the abundance of TAM subsets in CD8^low^TAM^hi^ and CD8^hi^TAM^hi^ tumors. **C)** Heatmap of biological pathway enrichment analysis (Reactome) based on DEGs comparing TAMs in CD8^low^TAM^hi^ and CD8^hi^TAM^hi^ tumors. Pathways absent in the data are marked in gray. **D)** Mean expression of selected IFN𝛾-induced genes comparing TAMs in CD8^low^TAM^hi^ and CD8^hi^TAM^hi^ tumors. ***p=0.001. **E)** Kaplan-Meier plot of 5-year overall survival related to the stromal density of GBP1, MCEMP1 and FOLR2 in MSS tumors. P-values are from log-rank test. **F)** Heatmap of biological pathway enrichment analysis (Reactome) based on DEGs comparing endothelial cells, epithelial (tumor) cells, stromal cells, and T/NK/ILCs in CD8^low^TAM^hi^ and CD8^hi^TAM^hi^ tumors. Pathways absent in the data are marked in gray. **G)** Mean expression of genes associated with antigen presentation, IFN type I, IFN type II, and chemokines expressed by tumor cells comparing CD8^low^TAM^hi^ and CD8^hi^TAM^hi^ tumors. ***p=0.001. **H)** Kaplan-Meier plot of 10-year overall survival related to IFN alpha/beta or gamma low and IFN alpha/beta or gamma high in MSS tumors. Upregulated genes in IFN alpha/beta signaling pathway (52 genes) and IFN gamma signaling pathway (58 genes) was used. Data from TCGA. P-values are from log-rank test.

The biological pathways enriched in TAMs within CD8^hi^TAM^hi^ tumors suggested a strong IFN-driven response, substantiated by the fact that many IFN type I and type II-inducible genes were significantly increased compared to CD8^low^TAM^hi^ tumors (Figure 7D). To examine whether this global polarization of TAMs had prognostic effect, we performed multiplex immunofluorescence staining of tissue microarrays (TMAs) from two large cohorts with a total of 1,720 CRC patients^20^ (Suppelementary Table 3) using the density of TAMs co-expressing CD68 and GBP1^59^ as a surrogate marker for an IFN-polarized TME. We designed two antibody panels that covered several TAM phenotypes/cell states. Antibodies to pan-macrophage markers CD68 and CD163 were included in both panels, together with DAPI and an antibody to cytokeratin to visualize nuclei and epithelial tumor cells, respectively. Panel 1 included antibodies to GBP1, MCEMP1 (preferentially expressed in proinflammatory TAMs), and CD206 (encoded by *MRC1*), often used to identify M2-polarized TAMs. Antibodies in panel 2 were specific for calprotectin (*S100A8/A9*) identifying recently recruited TAMs, ACP5 (preferentially expressed in *SPP1*+ TAMs), and *FOLR2*+ TAMs expressing a tissue resident phenotype (examples of staining results are given in Supplementary Figure 5A).

We focused on CRC patients with stage I-III tumors who underwent complete surgical removal of tumor tissue (R0 resection; n=1,370). As most TAMs are in the tumor stroma and because the tumor cells were variably present in the TMAs, we assessed the density of TAM phenotypes in the tumor stroma by subtracting the cytokeratin positive area (Supplementary Figure 5B). The density of each TAM phenotype was divided into three groups of similar size: high, intermediate, and low. We found that high density of *GBP1*+ TAMs had strong positive prognostic effects on both 5-year overall survival rate and 5-year relapse free survival in MSS tumors, but not in MSI tumors (Figure 7E and Supplementary Figure 5C). The density of *MCEMP*+ TAMs and *FOLR2*+ TAMs also showed significant, but modest, effect on prognosis of MSS tumors, whereas the density of the other TAM phenotypes (Calprotectin, CD206, and ACP5) had no prognostic effect (Supplementary Figure 5C). MSI tumors had significantly higher densities of TAMs expressing either MCEMP1, Calprotectin, GBP1, or ACP5, whereas *FOLR2*+ TAMs were more numerous in MSS tumors. The density of *CD206*+ TAMs showed no difference between the tumor types (Supplementary Figure 5C). Together, these results associated IFN-induced programming of TAMs with improved clinical outcome.

### The TME is enriched for immune-related programs in CD8^hi^TAM^hi^ tumors

Our findings suggested that either intrinsic properties of TAMs or different cell-extrinsic local cues polarized TAMs in the two tumor groups. If TAM polarization was driven by external signals, we hypothesized that this also would affect other cell types in the tumor tissue. To investigate this possibility, we determined the biological pathways of structural cells^60^ (epithelium (cancer cells), stromal and endothelial cells) and immune cells in the two tumor groups. Interestingly, this showed an enrichment for pathways resembling those in TAMs, including antigen presentation via class I and II/cross presentation, and IFN type I and II responses across all cell types in the CD8^hi^TAM^hi^ tumors, whereas pathways including TGFB-signaling, VEGFA-VEGFR2 pathway, PDGF signaling, and Th2 signaling were enriched CD8^low^TAM^hi^ tumors (Figure 7F). Several of the latter are involved in fibrosis and angiogenesis that are associated with worse prognosis in many cancer settings, including CRC^61–63^.

Interestingly to this end, differential abundance testing showed that several CAFs and tumor-associated capillary endothelial cells were more abundant CD8^low^TAM^hi^ tumors (Supplementary Figure 6A). Focusing on the malignant cell compartment we found increased expression of genes involved in antigen presentation, IFN type I ad IFN type II responses, as well as immune-attracting chemokines in CD8^hi^TAM^hi^ tumors (Figure 7G). These immune-related programs are very similar to the transcriptomic profile of malignant cells shown to be elevated in MMRd/MSI tumors^8,12^. To investigate the prognostic effect of IFN-induced TME we quantified the activation of DEGs in the IFN type I and type II signaling pathways (Reactome) upregulated in CD8^hi^TAM^hi^ tumors in publicly available RNAseq data from The Cancer Genome Atlas Program (TCGA). Interestingly, the overall survival in patients with MSS tumors was significantly better for both high levels IFN type I (p=1.7e-02) and type II (p=5.7e-04) in MSS tumors compared to tumors with low expression of these genes (Figure 7H).

Together, we find that structural cells including cancer cells and immune cells expressed an increased immune-related phenotype in CD8^hi^TAM^hi^ tumors and the transcriptomic profile of malignant cells in these tumors resembled that of MSI tumors.

### CD8 T cells are transcriptionally different in CD8^hi^TAM^hi^ and CD8^low^TAM^hi^ tumors

Differential abundance analysis confirmed that both *CXCL13+* and *CXCL13-* CD8 T cells were more abundant in CD8^hi^TAM^hi^ tumors (Supplementary Figure 6A). Moreover, both CD8 T cell subsets showed increased expression of genes related to immune activation (e.g. *IFNG*) and suppression (e.g. *PDCD1*) in the CD8^hi^TAM^hi^ group (Supplementary Figure 6B).

The chemokine receptor CXCR3 is important for recruitment of CD8 T cells and both CD8 T cells expressed significant levels of *CXCR3* (Figure 3C). In addition to TAMs and tumor cells (Figure 7D, G), the ligands for CXCR3 (*CXCL9, CXCL10*, and *CXCL11*), were also significantly increased in fibroblasts and endothelial cells in CD8^hi^TAM^hi^ tumors (Supplementary Figure 6C), suggesting that several cell types, besides TAMs, were involved in CD8 T cell recruitment.

Interestingly, Tregs, Th1-like, cm/naive T cells, and 𝛾/𝛿 T cells were more abundant in CD8^low^TAM^hi^ tumors, whereas IL17-like T cells, and NK cells were similar in both tumor types (Supplementary Figure 6A). This suggested that different mechanisms were involved in the accumulation of CD8 T cells and other T cell subsets and NK cells.

## DISCUSSION

CRC is a heterogeneous disease and various classification systems have proven useful for clinically stratifying patients, predicting prognosis, and guiding personalized treatment strategies^14,64^. Recently, we demonstrated that MSS tumors with high infiltration of T cells and TAMs exhibit significantly better clinical outcomes than MSS tumors with low T cell infiltration and high TAM density^20^. Here we used scRNA-seq, high resolution spatial transcriptomics, and multiplex immunohistochemistry on tumor tissue to uncover cellular and molecular mechanisms underlying this prognostic difference.

The prognostic impact of TAM infiltration in CRC has shown inconsistent results, with some studies linking high TAM density to improved outcomes and others associating it with poorer prognosis^65–70^. Our findings provide a potential explanation for these conflicting results by demonstrating that in MSS CD8^hi^TAM^hi^ tumors TAMs exhibit a polarization towards an IFN-induced phenotype. Moreover, the abundance of tumor-promoting *SPP1*+ TAMs was reduced in CD8^hi^TAM^hi^ tumors. Importantly, stratifying MSS CRC into CD8^hi^TAM^hi^ and CD8^low^TAM^hi^ tumors identified distinct co-regulated cellular programs across cell types of the TME. Both tumor, stromal, endothelial, and immune (T/NK/ILC) cells, and TAMs in CD8^hi^TAM^hi^ MSS tumors were enriched for biological pathways associated with anti-tumor immunity (e.g. antigen presentation, IFN type I and II responses), whereas CD8^low^TAM^hi^ tumors were enriched for pathways with tumor-promoting properties (e.g. TGFB-signaling, VEGFA-VEGFR2 pathway). These findings show similarities with a recent study by Bill et al. showing that CXCL9:SPP1 TAM polarity identified a network of cellular programs that control human cancers^32^.

We found that TAMs, cancer cells and other structural cells expressed higher levels of CXCR3-binding chemokines (*CXCL9, CXCL10, CXCL11*) in CD8^hi^TAM^hi^ tumors, which suggests that multiple cell types are important for the increased recruitment of tumor-infiltrating CD8 T cells. Importantly, *GBP1*+ TAMs and *CXCL13+* CD8 T cells were organized intratumorally in microniches (immune hubs) as suggested by others^32,45,46^. This situation mirrored a recent report by Elewaut et al uncovering a sequence of events of which IFN-induced cancer cells in concert with inflammatory TAMs (expressing *CXCL9* and *CXCL10*) drive intratumoral anti-tumor T-cell responses^46^. We find that these tumor-infiltrating *CXCL13+* CD8 T cells express high levels of inhibitory molecules (e.g. *PDCD1, CTLA4, LAG3*), which have been termed terminally exhausted^40^. However, this subset expressed the highest levels of *IFNG,* and a significant fraction were proliferating (*MKI67*+), indicating that they still were functionally active as suggested by others^41^.

In contrast to *CXCL13+* CD8 T cells, *CXCL13-GZMK+* CD8 T cells accumulated in TLS in the tumor stroma. Of particular interest we found close spatial relationship between mregDCs, *GMZK+* CD8 T cells, and *CXCL13+* CD4 T cells. These T cell subsets were only expressed in tumor tissue and several reports have shown that these phenotypes are tumor-reactive^35–41^. mregDCs expressed both high levels of T cell stimulatory and attracting molecules making them ideal to trigger anti-tumor T cell responses. Indeed, a recent report on hepatocellular carcinoma showed that the cellular triad of mregDCs, *GMZK+* CD8 T cells, and *CXCL13+* CD4 T cells was strongly associated with response to ICB. Similarly, expansion of *GZMK+* CD8 T cells was only detected in lung cancers that responded to anti-PD-1 treatments^42^. Together, these findings may partially explain why the occurrence of TLS is associated with response ICB^35,71^.

A subgroup of MSS CRC patients (27%) showed a pathological response to neoadjuvant immunotherapy. The strongest predictive response was infiltration of CD8+PD-1+ T cells^10^. Moreover, single-cell meta-analysis showed that CXCL13+ CD8 T cells correlated with favorable response to ICB^39^. It is therefore tempting to speculate that a classification system which stratify MSS tumors related to their densities of CD8 T cells and TAMs could aid the identification of CRC patients with MSS tumors that would benefit from ICB.

In conclusion, combining scRNA-seq and single cell spatial transcriptomics this study provides a comprehensive spatial transcriptomic atlas of the cellular and molecular landscape of CRC. Using a simple stratification system based on CD8 T cell and TAM densities, we uncover coordinated cellular programs across cell types within the TMEs of MSS tumors that are strongly associated with prognosis. Further investigation is warranted to determine whether this classification system could aid in identifying MSS CRC patients who may benefit from ICB ^10,72^.

## METHODS

### Data curation

Publicly available datasets from seven CRC studies were collected. Out of the seven curated datasets, three studies provided fastq files, while the remaining four studies had only raw count matrices available. The corresponding authors of these four studies were contacted in an attempt to require fastq files. However, due to lack of approval from the respective data access committees we were unable to obtain the fastq files. For these studies raw count matrices were used.

All studies contained scRNA-seq data from the primary CRC tumor, in addition to adjacent normal colon tissue except for Che et al.^22^ and Guo et al.^23^ who only included tumor data.

Sequencing data was downloaded from European Nucleotide Archive (Che et al.^22^ PRJNA725335, Guo et al.^23^ PRJNA779978), dbGaP (Pelka et al.^8^ phs002407), ArrayExpress database (Qian et al.^25^ E-MTAB-8107), GEO (Uhlitz et al.^26^ GSE166555) and European Genome-phenome Archive (Joanito et al.^12^ EGAD00001008555, EGAD00001008584, EGAD00001008585). Raw count matrices from Liu et al.^24^ were provided by the author upon request. Metadata files were downloaded from the supplementary files for each publication.

### Patient samples and processing of tissue

For single-cell RNA sequencing (scRNA-seq) analysis, colonic resections were acquired from patients (n=6, age range 62–78 years, three males, total of 6 samples) undergoing surgery for colorectal cancer at Akershus University Hospital (Ahus) and Oslo University Hospital (OUH). Clinical characteristics are detailed in Supplemental Table 4. Samples were processed as described in^21^. Briefly, samples of tumor tissue were digested for 60 minutes in RPMI [Lonza] supplemented with 10% FCS, 1% penicillin/streptomycin [Lonza], and 20 U/ml DNase I [Sigma-Aldrich] and 0.25 mg/ml Liberase TL [Roche]). Following filtration at 100-μm cells were stained on ice for 30 min with the following antibodies from BioLegend: CD45 BV510 (103138, clone 30-F11), HLA-DR PerCP/Cy5.5 (307630, clone L243) and CD14 APC (325608, clone HCD14). Stained cells were subjected to FACS sorting with a FACS Aria IIIu (BD Biosciences), and cellular suspensions (∼15,000 cells, with expected recovery of ∼7,500 cells) of sorted CD45+HLA-DR+CD14+ macrophages were loaded on the 10X Chromium Controller instrument (10X Genomics) according to the manufacturer’s protocol using 10X GEMCode proprietary technology. The Chromium Single Cell 3′ v2 Reagent kit (10X Genomics) was used to generate the cDNA and prepare the libraries, according to the manufacturer’s protocol. The libraries were then equimolarly pooled and sequenced on an Illumina NextSeq500 using HighOutput flow cells v2.5. A coverage of 400 million reads per sample was targeted to obtain 50,000 reads per cell. The raw data were then demultiplexed and processed with the Cell Ranger software (10X Genomics) v2.1.1.

For the TMA analysis the diagnostic biobank at OUH provided formalin-fixed and paraffin-embedded samples from the primary tumors of two separate cohorts of patients treated surgically for stage I–IV colorectal cancer (CRC) in a single hospital (n = 1720; Supplementary Table 3). These cohorts have been described in detail in previous publications^20,73^. They comprised 288 (17%) patients with a stage IV diagnosis of distant metastatic disease (three patients had missing data) and 1429 (83%) patients with a TNM stage I–III CRC (locoregional disease). Standard national guidelines were followed for the treatment and follow-up of the patients. From the patient’s medical records, clinicopathological data were registered into a standardized database.

As previously described^73^, TMAs were built from a single tissue core of the central tumor area of blocks chosen for representativeness by an experienced pathologist. Patients treated between 1993 and 2003 (n = 922) were included in Norwegian series 1 (NS1) and had 0.6 mm diameter cores. Patients treated between 2003 and 2012 (n = 798) were included in Norwegian series 2 (NS2), where TMAs with 1.0 mm diameter cores were used. Prior to this, scores were obtained for MSI status, BRAFV600E mutations, KRAS mutations^65,68–70^ and T cell and macrophage markers (CD3, CD8, and CD68)^20,73^.

### Preprocessing of fastq files and raw count matrices

Fastq files were processed using Cell Ranger version 7.0.0 with the default settings for alignment to the GRCh38 reference genome.

For raw count matrices, we verified the Cell Ranger version and reference genome to ensure they were compatible with other datasets.

### Quality control and integration

After sequence alignment, all samples were concatenated into a single object containing 1 522 014 cells, prior to performing the quality control (QC) using Scanpy^74^ (v1.9.8). Initially, we filtered the concatenated object by removing cells with fewer than 200 genes and genes expressed in fewer than 10 cells. Doublets were then removed from the object using SOLO^75^ (scvi-tools v1.1.1) using the default parameters. Additional filtration steps involved removal of outliers for “log1p_total_counts”, “log1p_n_genes_by_counts” and “pct_counts_in_top_20_genes” using a threshold of 5 median absolute deviations (MADs).

Normalization and identification of highly variable genes (n_top_genes set to 4000) were performed using Sanpy’s built-in functions. After QC, the concatenated object was reduced to 902 409 cells.

Integration of the concatenated object was performed using scVI^76^ (scvi-tools v1.1.1) with sample identifier as batch key, and sample identifier and study as categorical covariates. Additionally, percent mitochondrial reads and total counts were included as continuous covariates.

### Cell clustering, differential gene expression, and annotation

The integrated object was clustered using leiden clustering with a resolution of 0.1, which identified 11 distinct clusters. We used Scanpy’s built-in function for differential gene expression analysis, sc.tl.rank_genes_groups(), which employs Wilcoxon rank-sum test, with default parameters to identify differentially expressed genes between these clusters. Genes were considered as significantly differentially expressed if logFC > 0 and p-adjusted < 0.05. The significantly differentially expressed genes were used for cluster annotation, revealing major cell types such as epithelial, stromal, and immune cells. Each major cell type was then extracted and re-clustered to identify specific cell subsets.

### Multiplex fluorescence immunohistochemistry

Formalin-fixed and paraffin-embedded tissue sections of TMAs were cut in series at 4 μm and we used an automated Ventana Discovery ULTRA slide Stainer (Roche Diagnostics, Pleasanton, Calif.) for 7-color immunofluorescence staining. The stains were carried out with combination of primary antibodies and using a multiplex kit (NEL810001KT) together with Opal 620 (FP1495001KT, both from PerkinElmer/Akoya, Marlborough, MA, USA). Two different combinations consisting of 7 primary antibodies were assembled for staining of sequential sections. The primary antibodies used as a backbone in both setups were as follows: CD68 (M087601-2, clone PG-M1) visualized with Opal 690, CD163 (TA506382, clone OTI2B12) visualized with Opal 620, a cocktail of antibodies targeting the epithelial cancer cells (E-cadherin [clone 36, BD-biosciences, diluted 1:20,000], cytokeratin C-11 [Abcam, diluted 1:4000], cytokeratin Type I/II [Thermo Fisher Scientific, diluted 1:2000]) visualized with Opal 780, and included staining the cell nuclei with DAPI before mounting. In the first setup FOLR2 (MA5-26933, clone OTI4G6) was visualized with Opal Polaris 480, ACP5 (ab238033, clone rACP5/1070) visualized with Opal 520, and anti-human calprotectin from Calpro visualized with Opal 570. For the second combination GBP1 (LS-C786452-100, clone OTI1B2) was visualized with Opal Polaris 480, MCMP1 (NBP1-81253, polyclonal) visualized with Opal 520, and CD206 (91992S, clone E2L9N) visualized with Opal 570.

### Digital image analysis

Each image of a single TMA was processed using internally developed python scripts based upon the tiff-file library to read images and the cv2 library to process images. Snakemake was used to run all elements of the pipeline on all images in parallel^77^. Region segmentation was performed for each image where pixels were classified into “background”, “disregard”, “epithelium” and “stroma” using the Ilastik software^78^. “Disregard” is for example areas where the tissue is folded, “epithelium” has cytokeratin signal, “stroma” has DAPI signal but not “epithelium” and “background” is everything else. Pixel classification was done on images with the DAPI and cytokeratin channels that were clipped at the highest pixel value of each image. 38 images from both NS1 and NS2 and sigma2, sigma4 and sigma6 features were used to train a random forest classifier. The Ilastik command line tool was used to export one probability image for each label for all images in bulk.

For cell segmentation, the CD68 channel was extracted, clipped at an intensity of 45, and saved to a png file used to segment “CD68” and “background” using pixel classification in the Ilastik software. 11 images, using sigma2 and sigma5 features, were used for training the random forest classifier. The command line version of ilastik was used to batch process images giving one output image for probability of being “CD68” and one for probability of being “background”. The threshold for “CD68” probability image was created using a binary threshold with a lower threshold of 50, then the image was filtered using a morphology opening and a closing operation with a uniform kernel with size of 3 pixels. Pixel clusters smaller than 25 pixels were filtered out and contours were found using cv2.findContours(). Multiple centers were found for contours with an area larger than 450 pixels using cv2.kmeans() with n cluster set so that no contour was larger than 450 pixels. Contours were then split using the watershed algorithm using a halo size of 4 pixels. The sum of raw signal for each fluorescence channel as well as probabilities from region segmentation was then calculated for each contour and saved to a table for downstream processing.

PDFs were created showing the image used for region segmentation, the region segmentation, image used for CD68 segmentation as well as CD68 segmentation. All images were reviewed by looking through these PDFs and 132 images were excluded due to either bad CD68 segmentation (50 images), bad region segmentation (95 images) or both. A typical reason for exclusion is high CD68 background signal making segmentation unreliable for this image. Images with less than 1.5 % stromal area (73 images) or more than 80 % background (21 images) were filtered out, giving a total of 203 filtered out images.

Histograms of fluorescence signal for each antibody stain were used to determine thresholds for positive and negative cells. PDFs were then created of 50 random images to show CD68 signal, fluorescence signal and which cells were positive and negative for the specific antibody. These PDFs were used to verify that the fluorescence thresholds were appropriate.

### Survival analysis

CD68 contours that were not positive for stroma were filtered out and for each antibody stain the number of positive cells per mm2 of stroma was calculated. Clinical information was added to the data and negative control images, images from patients with TNM stage 4 and R-status not equal to zero were filtered out. log (antibody stain per mm2 stroma +1) was used to divide samples into quantiles. Non-numerical variables were converted into categorical variables and age was divided into 5 bins using the pandas.cut() function. The python package “lifelines” was used for survival analysis^79^. lifelines.CoxPHFitter() was used to perform Cox-regression and the lifelines.statistics.proportional_hazard_test() was used to test assumptions. No variable had a p-value from the proportional_hazard_test lower than 0.01. lifelines.KaplanMeierFitter() was used to generate KMP plots with p-values calculated with lifelines.statistics.logrank_test().

“rna_seq.augmented_star_gene_counts.tsv” files were downloaded from the Cancer Genome Atlas (TCGA) for 608 “adenomas and adenocarcinoma” patients with data from the colon. MSI/MSS status was downloaded from the supplementary of the paper describing the MANTIS program^80^. Samples from Primary tumor and with MSS status were selected. For each geneset, the average “tpm_unstranded” was calculated per patient. Patients were divided into two groups split at the 33 % percentile of this mean score. Survival data was downloaded^81^ and KMP plots and statistics was performed as above.

### Patient stratification into prognostic groups in scRNA-seq data

Patients were stratified into prognostic groups based on tumor data by calculating the median number of CD8+ T cells and tumor-associated macrophages (TAMs) per patient. Patients with fewer than 100 cells in total and/or fewer than 10 CD8+ T cells or TAMs were excluded from the analysis.

Patients with cell counts above the median were classified as CD8^hi^ or TAM^hi^, while patients with cell counts below the median were classified as CD8^low^ or TAM^low^. These classifications were combined to generate four prognostic groups based on CD8+ T cell and TAM abundance: CD8+ T cell^high^ TAM^high^, CD8+ T cell^low^ TAM^high^, CD8+ T cell^high^ TAM^low^, CD8+ T cell^low^ TAM^low^. Within these groups patients were further stratified based on MSI status, including only MSS patients in downstream analyses.

### Differential abundance analysis

We performed differential abundance analysis using Milo^27^ (Milopy v0.1.1), which generated a kNN graph for testing. For global-level analysis comparing cells from tumor to normal adjacent colonic tissue, we used the default parameters. At the local level, focusing on cell subsets comparing tumor to normal, a k value of 10 was used while keeping other parameters at their default settings. The same setup was used when comparing cell subsets in CD8^hi^TAM^hi^ MSS tumors to CD8^low^TAM^hi^ MSS tumors.

### Gene set enrichment analysis

To identify enriched pathways in tumor data, we applied PathfindR^82^ separately to each major cell lineage, comparing CD8^hi^TAM^hi^ MSS tumors to CD8^low^TAM^hi^ MSS tumors. PathfindR performs active-subnetwork-oriented pathway enrichment analysis, leveraging protein-protein interaction networks to enhance pathway discovery^82^. For this analysis, we selected Reactome as the reference database.

Differentially expressed genes between CD8^hi^TAM^hi^ MSS tumors to CD8^low^TAM^hi^ MSS tumors for each major cell lineage were used as input, and PathfindR was run under default settings.

### Visium HD

For the spatial transcriptomics analysis, we utilized three publicly available Visium HD samples from 10x Genomics^83^ in addition to four samples generated in our lab.

FFPE tissue sections (5 microns) were placed on Superfrost plus microscope slides (Epredia) and subjected to deparaffinization, H&E staining, imaging and decrosslinking according to the Demonstrated Protocol Visium CytAssist Spatial Gene Expression for FFPE (10x Genomics). Imaging was performed on an Olympus V200 slide scanner at 40x magnification. Probe hybridization, probe ligation, probe release, extension, library construction, and sequencing were performed according to the Visium HD Spatial Gene Expression Reagent Kits User Guide (10x Genomics). A CytAssist instrument was used to transfer transcriptomic probes from the glass slides to the Visium HD slides (6.5 mm capture area) according to the Visium CytAssist Spatial Gene Expression Reagent Kits User Guide (10x Genomics). Samples were sequenced in a NovaSeq 6000 S2 flow cell with 100 cycles. Reads were demultiplexed using the mkfastq command in Space Ranger (v3.0), and FASTQs were aligned to the human (GRCh38) reference with Space Ranger v3.0.

### Spatial clustering, cluster annotation, cell segmentation and spatial neighborhood analysis

Spatial clustering of bins from normal and tumor regions was performed using napari for area identification and bin annotation. Each bin was labeled according to its area of origin, and the spatial anndata object was subset to include only bins from normal or tumor regions for each sample. Per-sample objects were then integrated using scVI, followed by clustering and differential gene expression analysis as described above.

For cluster annotation transfer from single-cell to spatial data, we used our newly published Easydecon pipeline^28^. Briefly, the top 60 differentially expressed genes (DEGs), available on the Visium HD platform, for each major cell lineage were used to identify spatial bins most likely to contain the target cell type. For annotating dendritic cells (DCs), B cells, and plasma cells, manually selected markers were employed due to challenges in their identification from the spatial data. Epithelial cells were annotated using the 70th percentile, while the 95th percentile was used for other cell types (B cells, DCs, endothelial cells, macrophages, plasma cells, and T cells). Cell type annotations from were then transferred to the Visium HD data for spatial visualization using weighted Jaccard similarity (λ = 1), based on the top 60 DEGs for each cell type.

For cell segmentation, we applied Bin2Cell^29^, a method integrating morphological image segmentation and gene expression information for cell reconstruction. Morphological segmentation was performed using StarDist, a deep-learning-based tool for nuclei detection, leveraging its pretrained models to accurately identify cell nuclei. To refine segmentation, detected nuclei were expanded into neighboring unlabeled bins, ensuring a more complete cellular reconstruction. In parallel, gene expression data were utilized for secondary label identification, helping to infer cell identity in regions where nuclei were not captured or where cells had unusual shapes that evaded segmentation.

To assess spatial interactions between the transferred clusters, we performed spatial neighborhood analysis using Squidpy^34^ on Visium HD data. We first computed a spatial neighborhood graph, defining spatial relationships between spots. Next, we quantified neighborhood enrichment using, assessing whether specific clusters are spatially co-localized more than expected by chance.

## Data and code availability

Raw scRNA-seq data is deposited in EGA-archive (EGA accession number to be updated). Visum HD data and preprocessed scRNA-seq data are available via Zenodo (https://zenodo.org/uploads/15146453).

All source code is available on GitHub: https://github.com/JahnsenLab/CRC-atlas https://github.com/sinanugur/easydecon.

## ACKNOWLEDGEMENTS

We thank the Pathology and Surgery Departments at Oslo University Hospital and Ahus; Susanne Lorenz and Ana Heslanyeth Barragan Lid at the Genomic Core Facility, Oslo University Hospital; and members of the Jahnsen lab.

We sincerely thank Professors Ragnhild Lothe and Arild Nesbakken (both Oslo University Hospital) for access to the clinical database and the tissue microarrays from the patient cohorts Norwegian series 1 and 2.

The work was made possible through generous funding by South Eastern Norway Health Authority (no. 2022064 and 2022015), Norwegian Research Council (no. 315483), and the Norwegian Cancer Society (no. 90315, 223245, and 273170).

## AUTHOR CONTRIBUTIONS

Conceptualization: V.T.K., E.S.B., D.D., F.L.J.

Tissue processing and sequencing: K.T.H., A-C.R.B., U.M., E.S.B., V.T.K.

Patient recruitment and clinical input: M.A.M.

Perform multiplex immunochemistry: U.M., V.T.K., H.T.L., H.R.N.

Data analyses: V.T.K., S.U.U., D.D., U.M., E.S.B., F.L.J., H.T.L., K.R.V.

Atlas annotation: V.T.K., D.D., F.L.J. Writing – original draft: V.T.K., F.L.J.

Writing – review & editing: E.S.B., H.T.L., S.Y., D.D.

Supervision: F.L.J., D.D., E.S.B., S.Y.

## SUPPLEMENTARY TABLES

**Supplementary Table 1:** Overview of 60 differentially expressed genes (DEGs) per cluster and subcluster used for annotation.

**Supplementary Table 2:** Clinical characteristics of Visium HD samples.

**Supplementary Table 3:** Clinical characteristics of Tissue micro-array (TAM) samples.

**Supplementary Table 4:** Clinical characteristics of patients included in the scRNA-seq data.

---

**Supplementary Figure 1:**
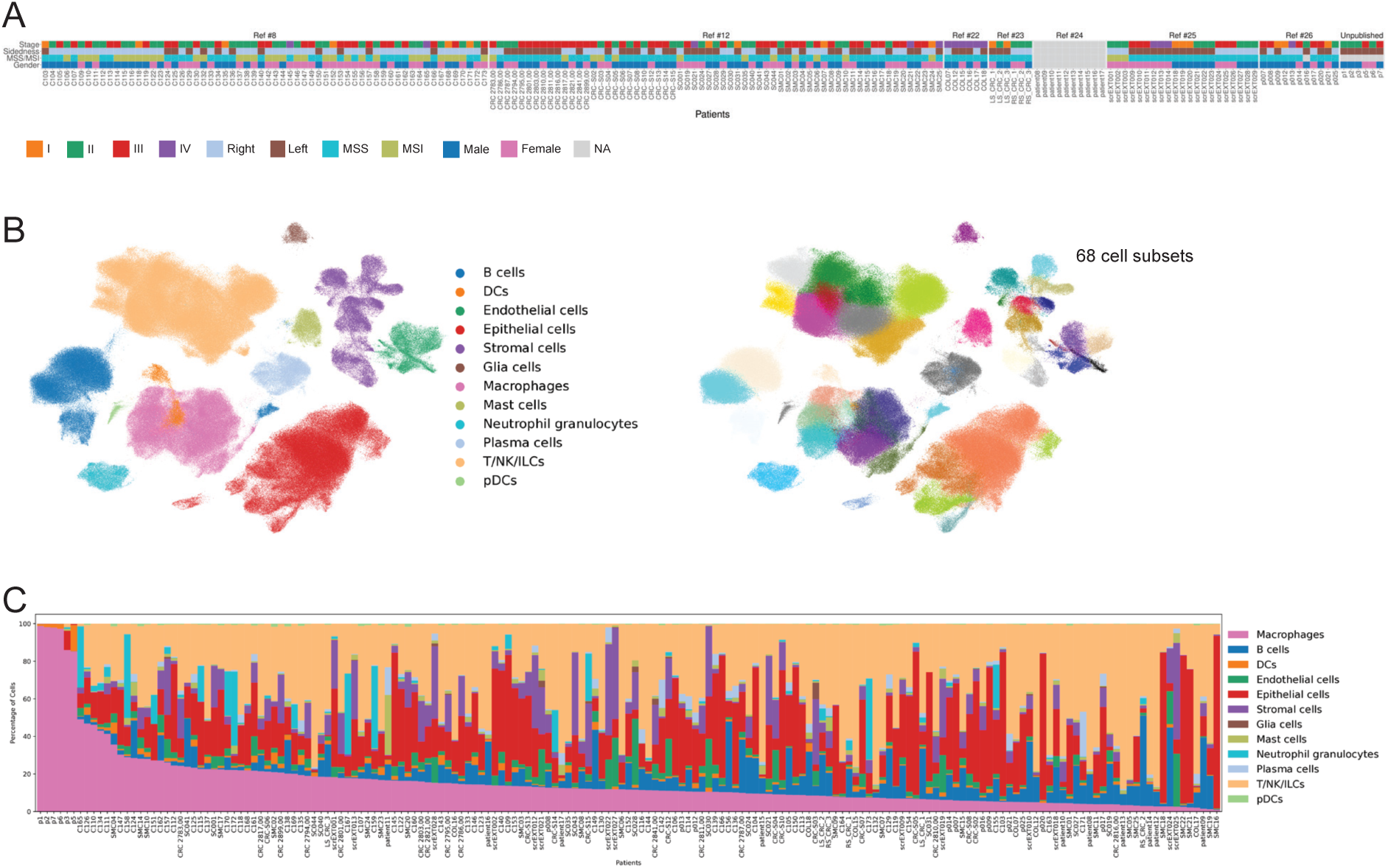
Patient characteristics and cell clustering. **A)** Clinical characteristics of CRC patients. **B)** UMAP for the major cell types combining scRNA-seq data from 185 CRC patients (left), and UMAP with fine-grained clustering of the cell types (right). **C)** Relative abundance of major cell types for individual CRC patients.

**Supplementary Figure 2:**
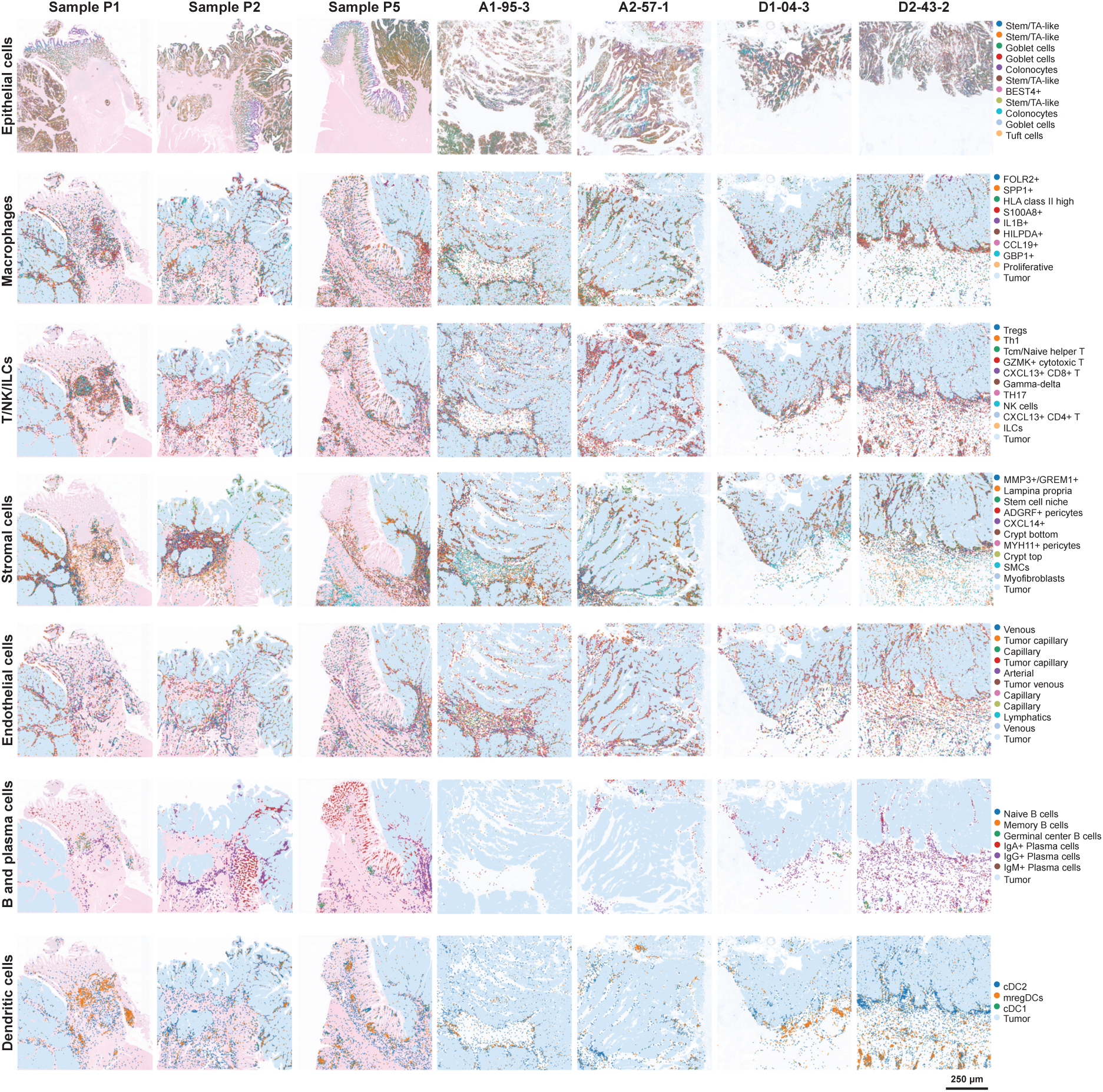
Spatial visualization combining scRNA-seq and Visium HD for all samples. Analysis of scRNA-seq data sets has been paired with Visium HD (Easydecon) performed on one FFPE section from seven CRC tumors. Each column (n=7) of images gives a nearly single-cell spatial resolution of 59 cell subsets on the same tissue section separated into major cell types (indicated to the left) for simplicity. Annotation of cell types are indicated (right). The three patient samples to the right are stained with H&E (Samples P1, P2, P3^83^), whereas the other four samples are stained with Hematoxylin only.

**Supplementary Figure 3:**
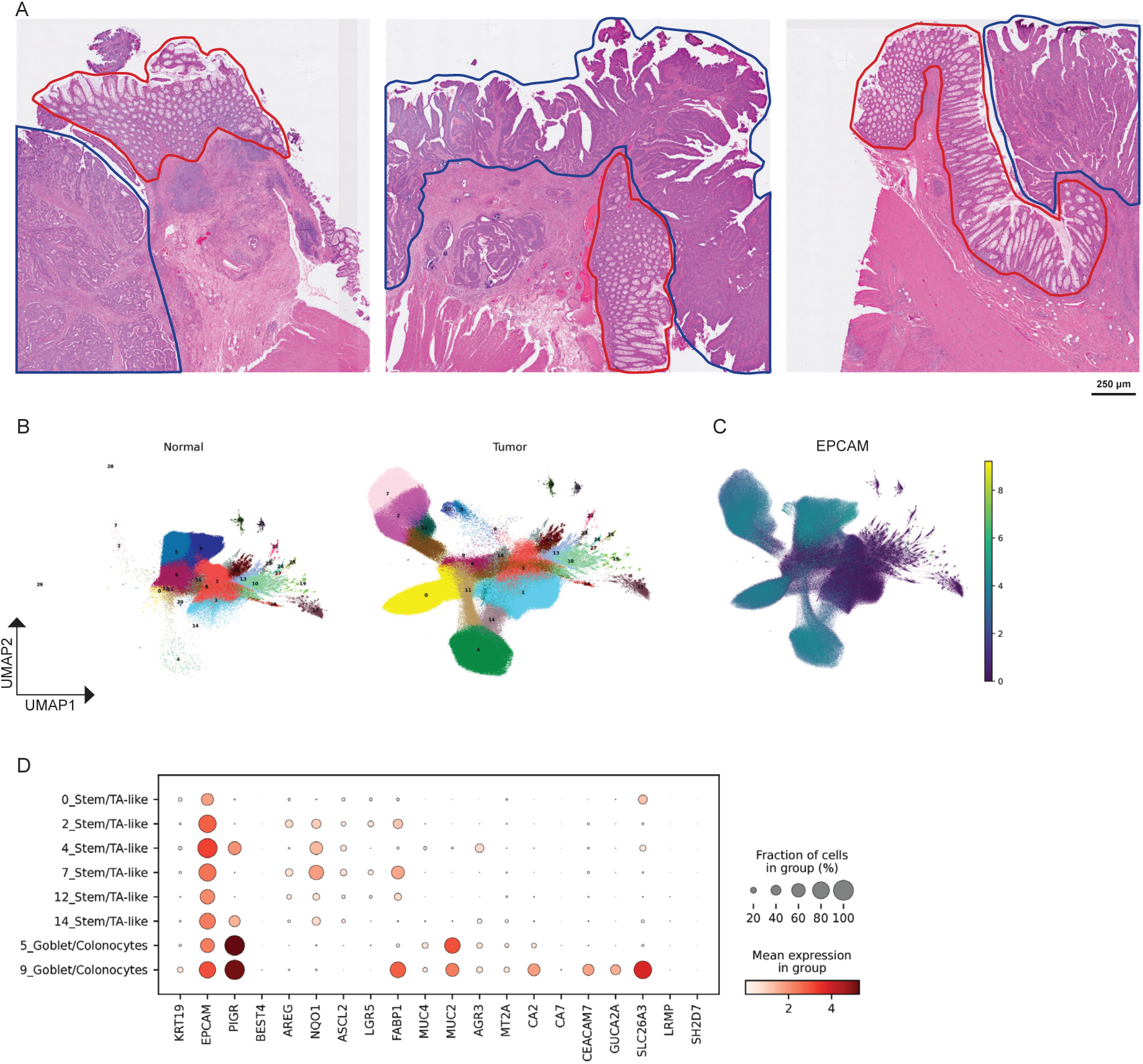
Visium HD performed as a stand-alone platform. **A)** Annotation of tumor tissue (blue) and normal epithelial tissue (red) on H&E-stained sections (samples P1, P2, P5^83^). **B)** UMAPs of bins derived from areas with normal epithelium and tumor tissue annotated in A. **C)** Feature plot of EPCAM expression in compiled data. **D)** Marker gene dot plot for epithelial cell subsets from *EPCAM+* clusters in B.

**Supplementary Figure 4:**
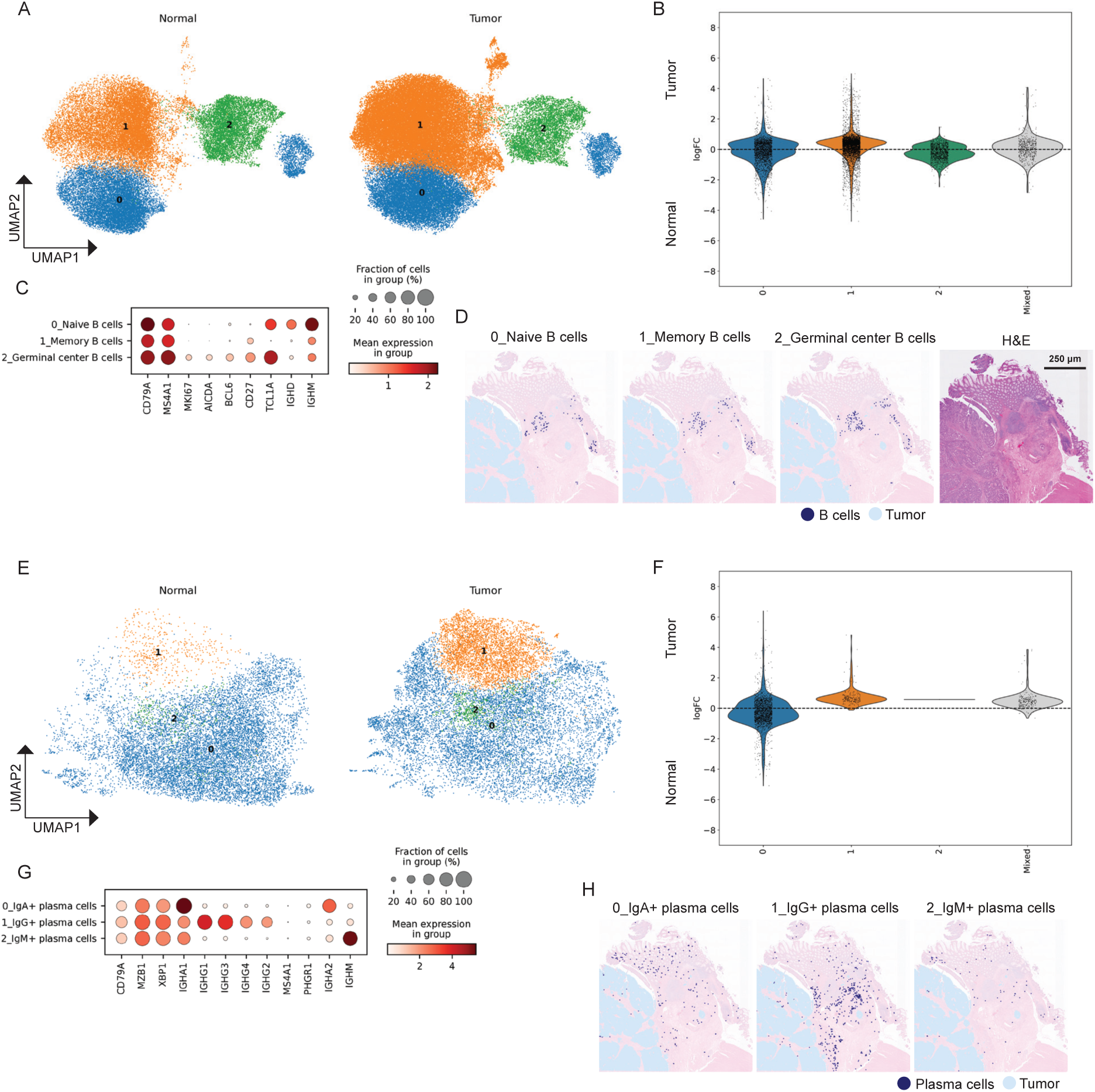
Analysis of B cells and plasma cells with scRNA-seq and Visium HD data. **A)** UMAPs of B cells from normal colon (left) and tumor tissue (right). **B)** Differential abundance analysis (Milo) comparing the abundance of B cells from normal colon and tumor tissue. **C)** Marker gene dot plot for B cell subsets from normal colon and tumor tissue with predicted annotations indicated. **D)** Spatial visualization of the annotated B cell subsets (A-C) combined with Visium HD (Easydecon) on FFPE tissue section from CRC (Sample P1^83^; each cell type is shown in separate images for simplicity). H&E staining (right). **E)** UMAPs of plasma cells from normal colon (left) and tumor tissue (right). **F)** Differential abundance analysis (Milo) comparing the abundance of plasma cells from normal colon and tumor tissue. **G)** Marker gene dot plot for cell subsets from normal colon and tumor tissue with predicted annotations indicated. **H)** Spatial visualization of the annotated plasma cell subsets (E-G) combined with Visium HD (Easydecon) on FFPE tissue section from CRC (Sample P1^83^; each cell type is shown in separate images for simplicity).

**Supplementary Figure 5:**
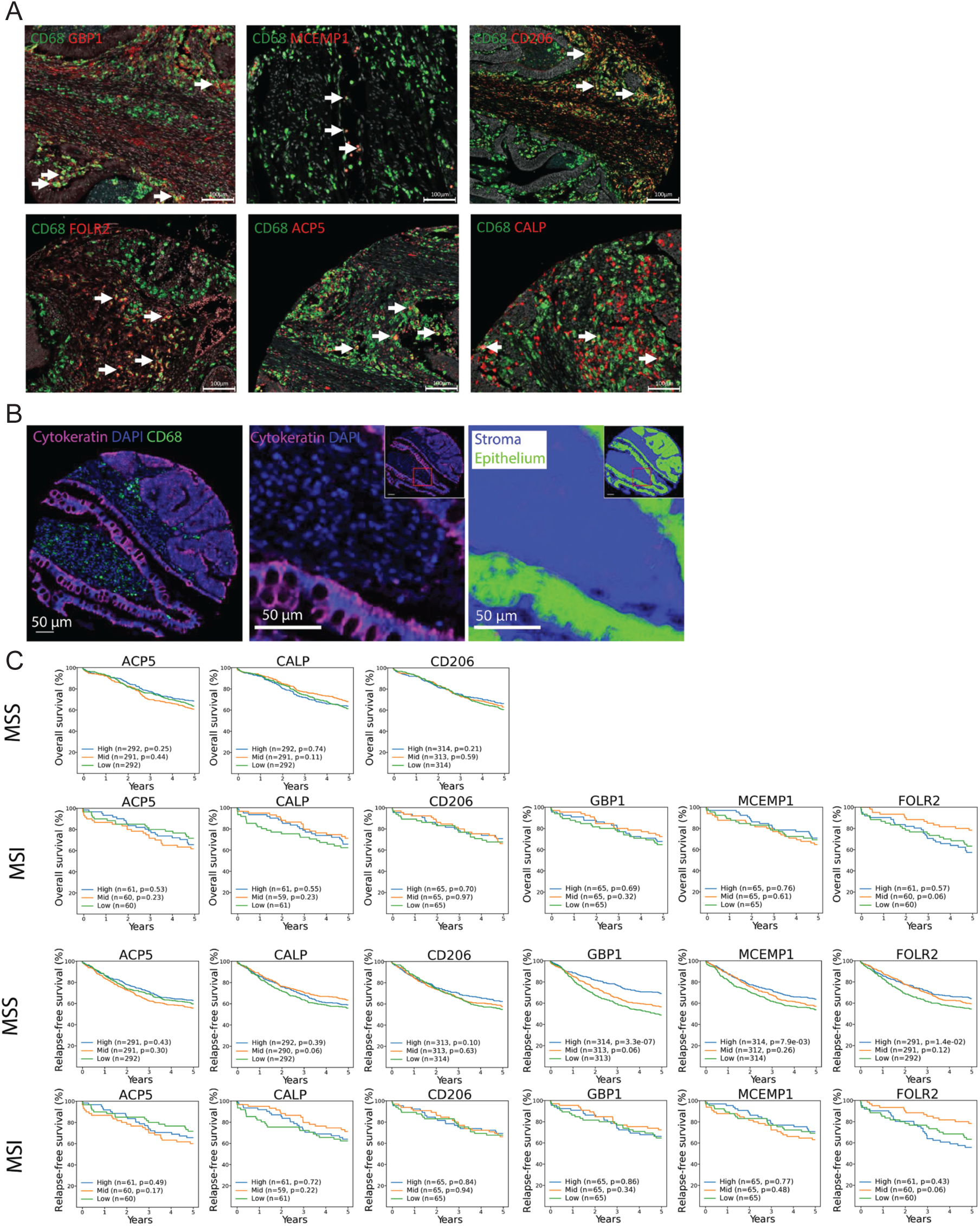
Analysis of TMAs from CRC tumors. **A)** Multicolor immunofluorescence stainings of FFPE TMA cores from tumor tissue with antibodies to CD68 (green) combined with either GBP1, MCEMP1, CD206, FOLR2, ACP5, CALP (calprotectin) (red) as indicated. White arrows show examples of TAMs positive for the indicated marker. Representative images are shown. **B)** Multicolor immunofluorescence stainings of FFPE TMA core from tumor tissue with antibodies to cytokeratin (magenta), CD68 (green), and DAPI to mark nuclei (blue). Representative image (right) of how cytokeratin positive area (green) was marked and subtracted to only count TAMs in the stromal area (blue). **C)** Kaplan-Meier plots of 5-year OS and RFS according to stromal density of TAM subsets as indicated for both MSS and MSI tumors. P-values are from log-rank test.

**Supplementary Figure 6:**
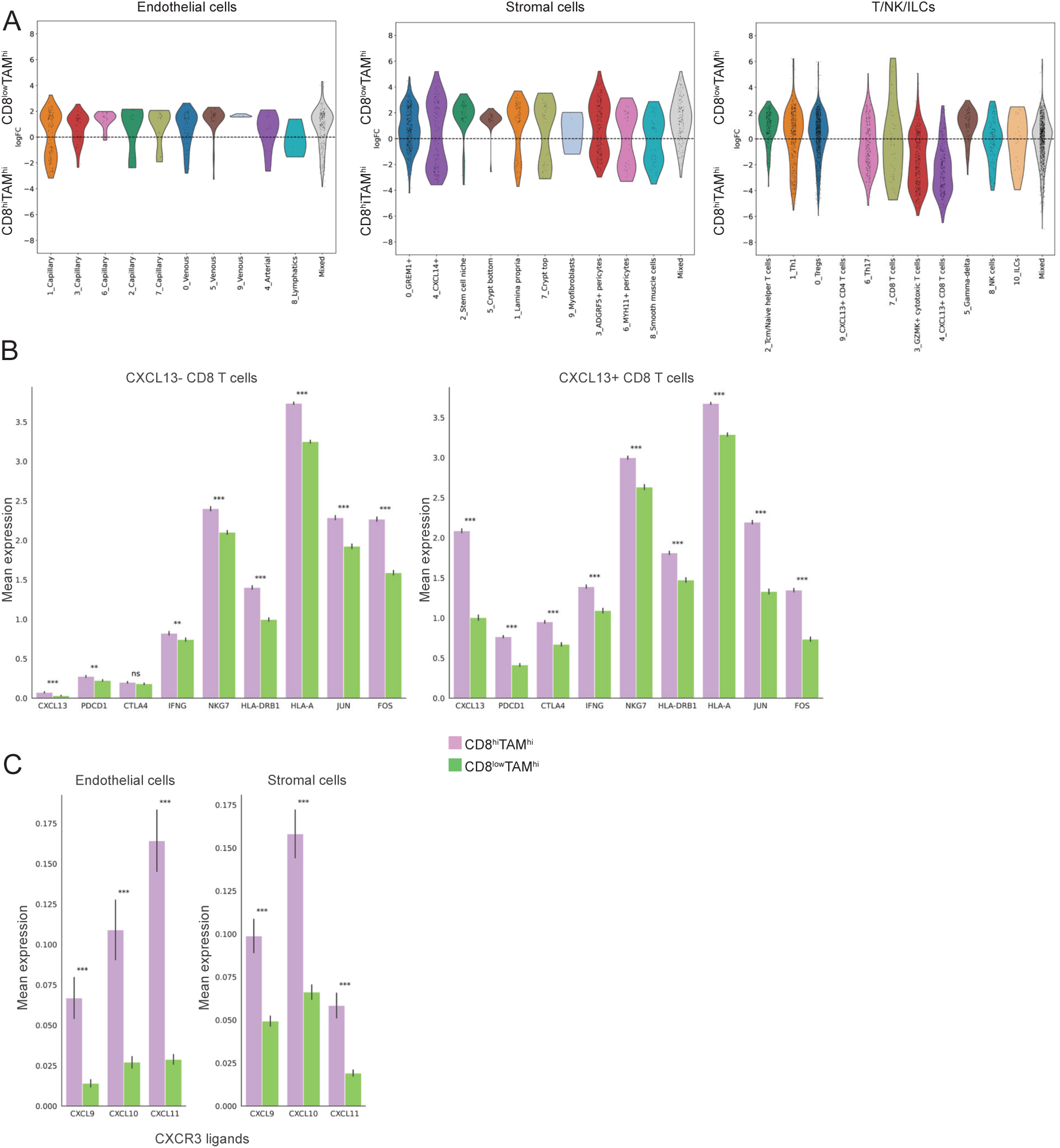
Transcriptional analysis of endothelial cells, stromal cells and T/NK/ILCs in CD8^low^TAM^hi^ and CD8^hi^TAM^hi^ MSS tumors. **A)** Differential abundance analysis (Milo) comparing the abundance of endothelial, stromal cells, and T/NK/ILC subsets in CD8^low^TAM^hi^ and CD8^hi^TAM^hi^ tumors. **B)** Mean expression of genes associated with immune activation and immune suppression expressed by *CXCL13+* and *CXCL13-* CD8 T cells in CD8^low^TAM^hi^ and CD8^hi^TAM^hi^ tumors. **p=0.01, ***p=0.001, ns=non-significant. **C)** Mean expression CXCR3-ligands *CXCL9, CXCL10*, and *CXCL11* by endothelial cells and stromal cells comparing CD8^low^TAM^hi^ and CD8^hi^TAM^hi^ tumors. ***p=0.001

