## Supplementary Table 3 for "Single cell spatial transcriptomics identifies coordinated cellular programs associated with good prognosis in microsatellite stable colorectal cancer"

| | | Norwegian series 1 (1993-2003) | Norwegian series 2 (2003-2012) | Total |
|---|---|---|---|---|
| Total patients, n | | 922 | 798 | 1,720 |
| Age | Median (range) | 73 (29–94) | 72 (27–97) | 72 (27 – 97) |
| Sex | Female | 485 (53%) | 407 (51%) | 892 (52%) |
| | Male | 437 (47%) | 391 (49%) | 828 (48%) |
| TNM stage | I | 137 (15%) | 167 (21%) | 304 (18%) |
| | II | 381 (41%) | 288 (36%) | 669 (39%) |
| | III | 242 (26%) | 214 (27%) | 456 (27%) |
| | IV | 159 (17%) | 129 (16%) | 288 (16) |
| | NA | 3 | - | 3 |
| pT – Tumor invasion | 1 | 37 (4%) | 40 (5%) | 77 (4%) |
| | 2 | 127 (14%) | 165 (21%) | 292 (17%) |
| | 3 | 662 (72%) | 515 (65%) | 1,177 (69%) |
| | 4 | 96 (10%) | 72 (9%) | 168 (10%) |
| | NA | 0 | 6 | 6 |
| pN – Nodal Involvement | 0 | 563 (62%) | 491 (62%) | 1,054 (62%) |
| | 1 | 250 (27%) | 175 (22%) | 425 (25%) |
| | 2 | 99 (11%) | 129 (16%) | 228 (13%) |
| | NA | 10 | 3 | 13 |
| Residual tumor (R) status | R0 | 719 (78%) | 651 (82%) | 1,370 (80%) |
| | R1 | 36 (4%) | 19 (2%) | 55 (3%) |
| | R2 | 167 (18%) | 128 (16%) | 295 (17%) |
| Tumor location | Right colon | 365 (40%) | 327 (41%) | 692 (40%) |
| | Left colon | 301 (33%) | 239 (30%) | 540 (31%) |
| | Rectum | 231 (25%) | 218 (27%) | 449 (26%) |
| | Synchronous | 25 (3%) | 14 (2%) | 39 (2%) |
| MSI status | MSI | 128 (15%) | 120 (16%) | 248 (16%) |
| | MSS | 712 (85%) | 638 (84%) | 1,350 (84%) |
| | NA | 82 | 40 | 122 |
| $BRAF^{V600E}$ mutational status | Wild-type | 714 (85%) | 637 (84%) | 1,351 (84%) |
| | Mutated | 127 (15%) | 122 (16%) | 249 (16%) |
| | NA | 81 | 39 | 120 |
| KRAS mutational status | Wild-type | 463 (69%) | 238 (69%) | 701 (69%) |
| | Mutated | 204 (31%) | 106 (31%) | 310 (31%) |
| | NA | 255 | 454 | 709 |
| Adjuvant chemotherapy | No | 806 (87%) | 609 (79%) | 1,415 (84%) |
| | Yes | 116 (13%) | 162 (21%) | 278 (16%) |
| | NA | 0 | 27 | 27 |
